# Proteomic profiling of the oncogenic septin 9 reveals isoform-specific interactions in breast cancer cells

**DOI:** 10.1101/566513

**Authors:** Louis Devlin, George Perkins, Jonathan R. Bowen, Cristina Montagna, Elias T. Spiliotis

**Author notes:** Corresponding author: Elias T. Spiliotis, Department of Biology, Drexel University, PISB 423, 3245 Chestnut St, Philadelphia, PA 19104, USA.

## Abstract

Septins are a family of multimeric GTP-binding proteins, which are abnormally expressed in cancer. Septin 9 *(SEPT9)* is an essential and ubiquitously expressed septin with multiple isoforms, which have differential expression patterns and effects in breast cancer cells. It is unknown, however, if SEPT9 isoforms associate with different molecular networks and functions. Here, we performed a proteomic screen in MCF-7 breast cancer cells to identify the interactome of GFP-SEPT9 isoforms 1, 4 and 5, which vary significantly in their N-terminal extensions. While all three isoforms associated with SEPT2 and SEPT7, the truncated SEPT9_i4 and SEPT9_i5 interacted with septins of the SEPT6 group more promiscuously than SEPT9_i1, which bound predominately SEPT8. Spatial mapping and functional clustering of non-septin partners showed isoform-specific differences in interactions with proteins of distinct subcellular organelles (e.g., nuclei, centrosomes, cilia) and functions such as cell signaling and ubiquitination. Notably, the interactome of the full length SEPT9_i1 was more enriched in cytoskeletal regulators, while the truncated SEPT9_i4 and SEPT9_i5 exhibited preferential and isoform-specific interactions with nuclear, signaling and ubiquitinating proteins. These data provide evidence for isoform-specific interactions, which arise from truncations in the N-terminal extensions of SEPT9, and point to novel roles in the pathogenesis of breast cancer.

## Introduction

Septins are a large family of GTP-binding proteins that control the intracellular localization of cytoskeletal, membrane and cytosolic proteins (1–5). Septins interact with one another via their GTP-binding domains (G-domains), forming homomeric and heteromeric complexes which assemble into higher-order filamentous structures (6, 7). Septins associate with the actin and microtubule cytoskeleton as well as cell membranes, and function in a variety of cellular processes including cell division and migration, intracellular membrane traffic, cell proliferation and apoptosis (5, 8, 9).

Based on sequence similarity, mammalian septins are classified into four groups named after SEPT2 (SEPT1, SEPT2, SEPT4, SEPT5), SEPT6 (SEPT6, SEPT8, SEPT10, SEPT11, SEPT14), SEPT7 and SEPT3 (SEPT3, SEPT9, SEPT12) (10–12). Septin paralogs from each of these four groups form complexes of various sizes and composition such as the palindromic hexamers and octamers of SEPT2/6/7 and SEPT2/6/7/9, respectively, which are considered as the minimal units of septin filaments (13–15). While the precise identity and diversity of septin complexes eludes our knowledge, septin-septin interactions are influenced by GTP binding and hydrolysis, the relative abundance of septin paralogs and isoforms, which can vary between cell types, and the presence of domains (e.g., N-terminal extensions) that may interfere with the G-domain binding interface (9, 16–19). Importantly, the localization and function of septin complexes appear to depend on the properties and binding partners of their individual subunits (20–22).

Septin 9 (SEPT9) is a ubiquitously expressed septin, which is essential for embryonic development; SEPT9 knock-out mice die early in gestation (23). A salient feature of SEPT9 is the presence of a long N-terminal extension (NTE), which is unique among septins. Owing to alternative translation start and splicing sites, there are at least five different SEPT9 isoforms (SEPT9_i1 to _i5) with NTEs that vary in length and amino acid sequence (24). The NTE of SEPT9 is structurally disordered and its full-length amino acid sequence comprises a domain (1–148) of basic isolectric point, which contains microtubule- and actin-binding sites, and an acidic proline-rich domain (aa 148-256) (25, 26). SEPT9 isoforms associate with the SEPT7 subunits of the hetero-hexameric SEPT7-SEPT6-SEPT2-SEPT2-SEPT6-SEPT7 complex, but longer SEPT9 isoforms might be preferred over shorter ones (14, 15, 20). Interestingly, in some cell types and processes, SEPT9 proteins appear to localize and function independently of SEPT2/6/7 (19, 27, 28).

SEPT9 was among the first septins implicated in cancer as the *SEPT9* gene was mapped to the loss of heterozygosity region of chromosome 17 (17q25), which is frequently deleted in breast and ovarian cancers (29, 30). The *SEPT9* gene was also found as a fusion partner of mixed lineage leukemia (MLL) gene and a preferred site of integration for a murine retrovirus that causes T-cell lymphomas (31, 32). In breast carcinomas and mouse models of breast cancer, the *SEPT9* gene is amplified (multiple gene copies) and the expression levels of certain SEPT9 variants are markedly increased (33, 34). Over-expression of SEPT9 isoforms occurs in ~30% of human breast cancer cases and correlates with poor prognosis and resistance to anti-cancer agents that target microtubules (34–38). Notably, changes in the methylation status of the *SEPT9* gene have been utilized for early detection of colorectal cancer (39, 40). Several studies have shown that over-expression of SEPT9 isoforms has differential effects on the migratory properties of breast cancer cells and have linked SEPT9_i1 to mechanisms of angiogenesis and cell proliferation (41–45). Despite these findings, we have a very poor understanding of the roles that different SEPT9 isoforms play in the development and metastasis of breast cancer.

Unbiased shotgun proteomics, which is based on liquid chromatography coupled to tandem mass spectrometry (LC-MS/MS), has been an effective approach in elucidating the molecular networks and complexes of cellular proteins (46, 47). Combined with bioinformatics, discovery-based proteomics can provide a comprehensive map of the cellular components, pathways and functions of a protein and its binding partners (48). Yeast hybrid screens have used mammalian septin paralogs as baits to identify potential binding partners (49, 50), but there is a dearth of proteomic data on septin complexes isolated from mammalian cells. Owing to the lack of isoform-specific antibodies, comparative proteomics for septin isoforms are challenging and septin isoform-specific interactomes are underexplored. This is a key challenge for SEPT9, which has a multitude of isoforms that may have differential roles in breast cancer. Here, we used discovery-based shotgun proteomics in MCF7 breast cancer cells, which were stable transfected with GFP-tagged SEPT9 isoforms 1, 4 and 5. We chose these isoforms because their amino acid sequences have the least overlap among all SEPT9 isoforms, and therefore, may have divergent interactomes and functions. Proteomic profiling revealed isoform-specific differences in the interaction with septin paralogs, and non-septin proteins of distinct subcellular localization (e.g., nucleus, centrosome, cilia) and functions including cell signaling and ubiquitin-mediated degradation. Collectively, these data suggest a functional specialization for isoforms of SEPT9, which arises from differences in the length and sequence of the N-terminal extensions of their GTP-binding domains.

## Materials and Methods

### Cell culture

MCF7 cells stably expressing GFP-SEPT9_i1, GFP-SEPT9_i4 and GFP-SEPT9_i5 (41) were cultured in Dulbecco’s Modified Eagle’s Medium with high glucose (4500 mg/L), L-glutamine and sodium pyruvate plus 10% fetal bovine serum and the following antibiotics: kanamycin sulfate, penicillin G and streptomycin sulfate. Cells were grown to confluency in 10 cm dishes at 37°C supplemented with 5% CO2.

### Fluorescence microscopy

MCF-7 cells expressing GFP-tagged isoforms of SEPT9 were fixed with warm PHEM buffer (60 mM Pipes-KOH, pH 6.9, 25 mM Hepes, 10 mM EGTA and 1 mM MgCl2) containing 4% PFA (EM Sciences) and 0.1% Triton X-100 and stained with a mouse antibody to α-tubulin (DM1α; SIGMA) and a secondary donkey DyLight 594-conjugated F(ab’)2 mouse IgGs (Jackson ImmunoResearch Laboratories, Inc) and Rhodamine-phalloidin (Cytoskeleton). Samples were mounted in Vectashield (Vector Laboratories) and imaged with an Olympus Fluoview FV1000 confocal laser scanning microscope with a Plan Apochromat 60X/1.35 NA objective and 488 nm, 543 nm and 635 nm laser lines. Three-dimensional stacks were collected at 0.2 μm-step size. Images were exported to the Slidebook 4.2 software and maximum intensity projections of a selected range of optical planes was generated before importing into Adobe Photoshop, where images were adjusted to 300 dpi before cropping for inclusion in the manuscript’s figures, which were made in Adobe Illustrator.

### Immunoprecipitation (IP)

Cells were washed twice with 2mL PBS at 4°C. 400 μL of IP lysis buffer (0.025M Tris, 0.15M NaCl, 0.001M EDTA, 1% NP-40, 5% glycerol; pH 7.4) supplemented with 2X Halt™ Protease Inhibitor Cocktail (ThermoFisher Scientific) was added to each plate and incubated on a rocker for 10 minutes at 4°C. Cells were scraped and lysate was transferred to a microcentrifuge collection tube and incubated end-over-end for 20 minutes at 4°C. Microcentrifuge collection tubes were then centrifuged at 16,000 xg for 10 minutes and supernatant was collected and frozen at −70°C until use.

Immunoprecipitations (IPs) were performed using Pierce Co-IP Kit (ThermoFisher Scientific) according to manufacturer’s instructions. Briefly, goat polyclonal anti-GFP antibody (Abcam ab6673) was coupled to AminoLink Plus Coupling Resin (10 μg/50 μL) using sodium cyanoborohydride. Antibody-coupled resin was incubated with analyte-containing lysate overnight with gentle rocking at 4°C. Five independent experiments for each isoform were performed. In each experiment, three 10 cm plates of equivalent confluency were individually lysed and IPs were performed in each of three lysates. The eluted complexes from all thee IPs were pooled into a single vial and dried at ambient temperature overnight using the SPD1010 SpeedVac system (Thermo Scientific).

### In-solution Protein Digestion

Vacuum-dried protein pellets were reconstituted in 0.1% (w/v) *RapiGest* SF Surfactant (Waters) in 50 mM ammonium bicarbonate. Samples were reduced with 5mM dithiothreitol (DTT; ThermoFisher Scientific; cat #: 20291) at 60°C for 30 minutes. Samples were then cooled to room temperature and free cysteines were alkylated with 15 mM iodoacetamide (ThermoFisher Scientific) in the dark at room temperature for 30 minutes. Samples were digested at 37°C overnight in 10 μg MS-grade Trypsin Gold (Promega). Sequencing grade trifluoroacetic acid (TFA; ThermoFisher Scientific) was added to the digested proteins to a final 0.5% concentration and incubated at 37°C for 45 minutes. Samples were centrifuged at 13,000 RPM for 10 minutes after which the supernatant was transferred to an HPLC low volume sample vial.

### LC-MS/MS Protein Identification

Each sample was analyzed using online liquid chromatography (Accela pump and autosampler, Thermo, Inc.) coupled to an LTQ-XL linear ion trap mass spectrometer. Samples were loaded onto an UPLC Peptide CSH C18 Column, 130 Å, 1.7 μm, 1 mm x 100 mm (Waters) maintained at 40°C. Solvent A and B were 0.1% formic acid in water and 0.1% formic acid in acetonitrile, respectively (gradient: 98%A/2%B for 59.5 minutes, 50%A/50%B for 0.5 minutes, 98%A/2%B for 10 minutes). The flow rate was set to 100 μL/min and the injection volumes were 10 μL/sample. The samples were ionized using ESI and fragmented using CID. Data were acquired in DDA mode. The mass spectrometer was run in positive ion mode using a source voltage of 3.50 kV, a capillary voltage of 40 V and a capillary temperature of 325°C.

Mass spectra extraction and deconvolution were performed using PEAKS Studio (Bioinformatics Solutions Inc. v8.5). Mass spectra were searched against the *Homo sapiens* database (UniprotKB) using a precursor mass 1.0 Da (monoisotopic mass) and a fragment ion mass 0.5 Da. Non-specific cleavages were allowed at both ends of the peptide and the number of allowed missed cleavages was set to 5. Specified variable modifications included: carbamidomethylation, deamidation (NQ) and oxidation (M). False positives were eliminated through a systematic four-tier process, which excluded proteins based on the following criteria: i) proteins that did not satisfy a P-value (probability of false identification) score of −10lgP ≥ 15 in peptide-to-spectrum and peptide-to-protein sequence matching by the PEAKS Bioinformatics software (see Supplementary Information, Table S1 for all protein hits with −10lgP ≥ 15); ii) proteins identified in IPs from non-transfected (control) cells; iii) proteins that were not identified in at least two out of five independent IPs; iv) proteins with a score of > 150 in the contaminant repository for affinity purification (CRAPome; www.crapome.org) (51). Based on known caveats of the “two peptide” rule which often results in increased false discovery rates (52), protein hits with single peptide matches were retained if they met a peptide-to-spectrum score of −10lgP ≥ 15 and peptide-to-protein matches were unique to a protein group, which was further verified by manual blasting of peptide sequences against the UniProt database. For all single peptide hits, annotated MS/MS spectra with precursor mass, charge and mass error, retention times, number of spectral matches and −10lgP scores are provided in Supplementary Information, Table S2.

### Bioinformatic analyses & data graphing

Protein hits were cross-checked against the Biological General Repository for Interaction Datasets (BioGRID; https://thebiogrid.org), septin interactomes (49, 54) and reviews of septin interactions (22), which were used for *in litero* verification of interactions (asterisks in Figure 2) and to generate Table I. Non-septin protein hits were spatially mapped by assigning each individual protein into one or more subcellular organelles and structures based on knowledge of protein localization in databases such as the human protein atlas (www.proteinatlas.org), GeneCards (www.genecards.org), UniProt (www.uniprot.org), COMPARTMENTS (https://compartments.jensenlab.org) and the Gene Ontology Consortium (www.geneontology.org). Subcellular localizations of protein were further corroborated and researched with pubmed (https://www.ncbi.nlm.nih.gov/pubmed/) queries. A combination of these databases and pubmed searches were used for clustering protein hits into functional categories.

**Figure 1.**
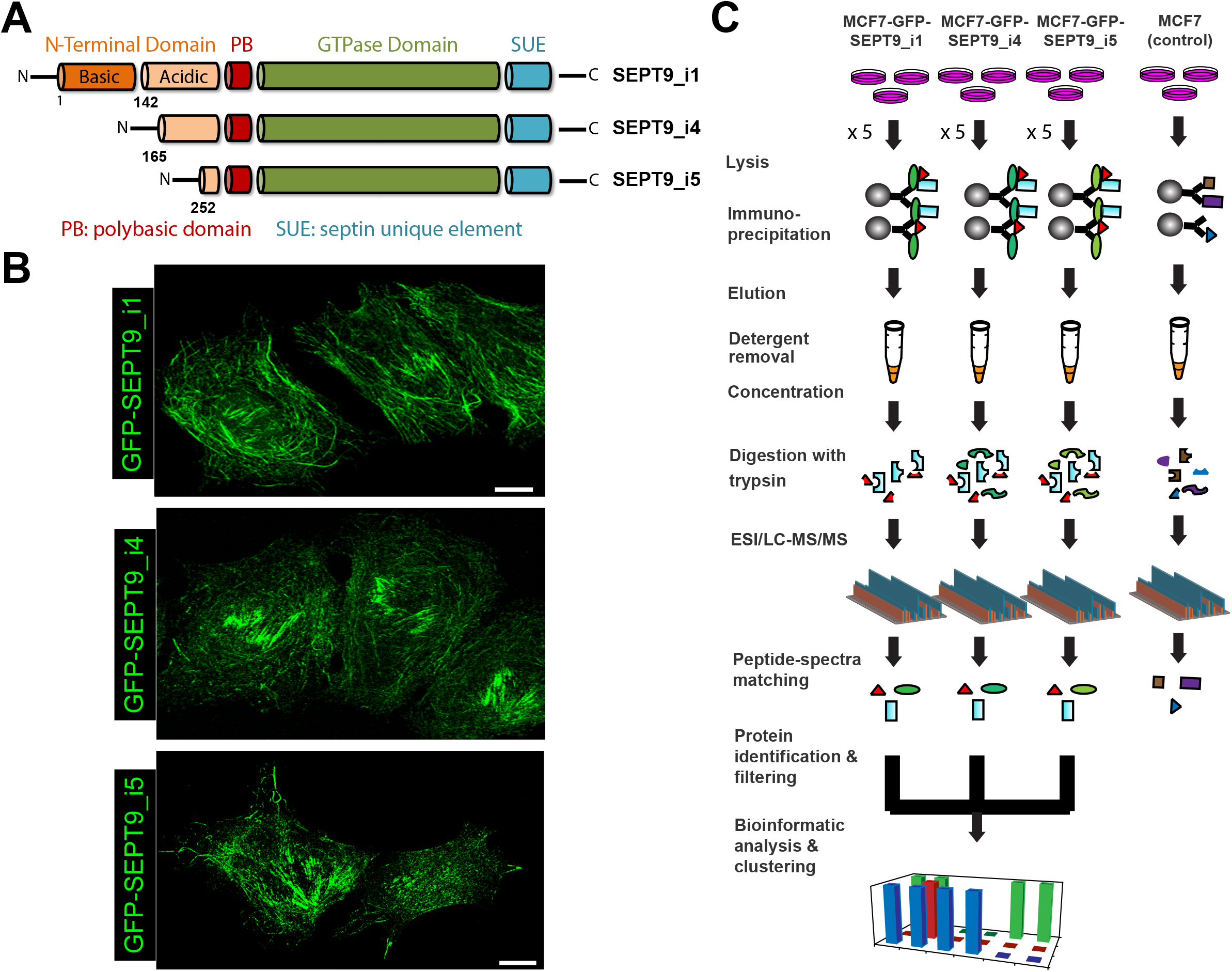
Proteomic screen for interactors of SEPT9 isoform 1, 4 and 5 in MCF-7 breast cancer cells. (A) Schematic depicting the lengths and major domains of the human SEPT9_i1, SEPT9_i4 and SEPT9_i5, which vary in the length at their N-terminal extensions. SEPT9_i4 lacks the N-terminal 164 amino acids of SEPT9_i1, and SEPT9_i5 lacks the N-terminal 251 and 87 amino acids of SEPT9_i1 and SEPT9_i4, respectively. (B) Confocal microscopy images of MCF-7 cells that stably express GFP-SEPT9_i1, GFP-SEPT9_i4 and GFP-SEPT9_i5. (C) Schematic of the proteomic screen which was performed five independent times with MCF-7 cell lines expressing GFP-SEPT9_i1, GFP-SEPT9_i4 and GFP-SEPT9_i5 as well as nontransfected MCF-7 cells (negative control). In each independent experiment, three sub-confluent plates with MCF-7 cells were lysed and immunoprecipitations against GFP were performed in each of the three lysates. Eluants were pooled, vacuum dried and digested with trypsin after detergent removal for injection into a liquid chromatography (LC) unit coupled to an electrospray ionization mass spectrometer (ESI-MS). Mass spectra were extracted and protein identities were derived by searches against the UniprotKB database. Data were filtered by applying quantitative thresholds to correct for non-specific interactions and contaminants, low-quality peptide-spectra and peptide-protein matches as well as low reproducibility. Data were analyzed for overlapping and unique binding partners, and non-septin binding partners were binned into subcellular organelle/structure and functional clusters.

**Figure 2.**
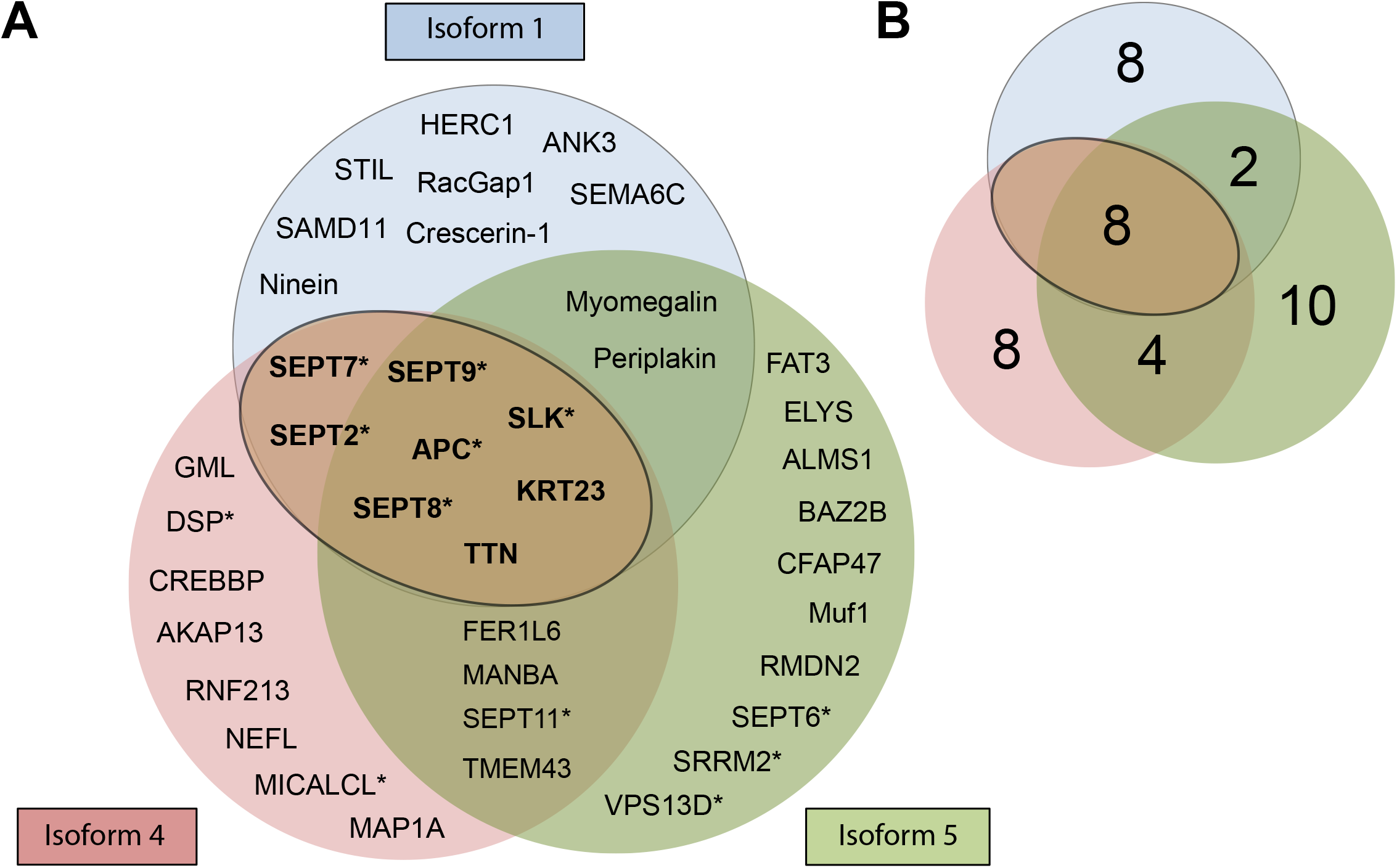
SEPT9 isoforms 1, 4 and 5 have overlapping and distinct interactors in MCF-7 breast cancer cells. Venn diagrams depicts the names (A) and cumulative numbers (B) of distinct and overlapping interactors of GFP-SEPT9 isoforms 1 (blue), 4 (pink) and 5 (green) in MCF-7 breast cancer cells. The overlapping interactors (bold letters) between isoforms 1, 4 and 5 are the same with the overlapping interactors of isoforms 1 and 4, and are shown in the outlined brown oval. Asterisks denote previously reported interactions.

**Table I:**
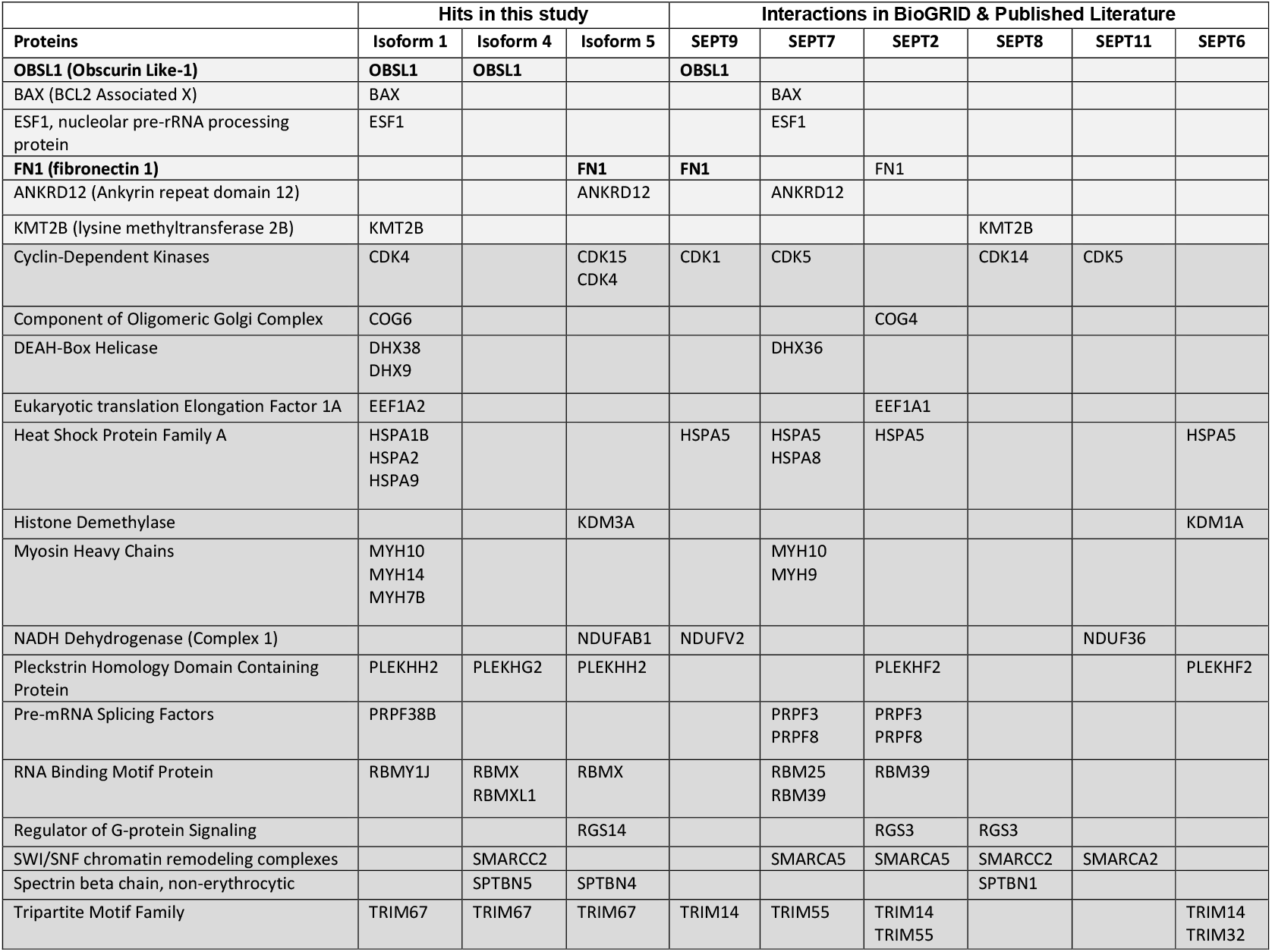
Septin-interacting proteins that were pulled down but did not meet threshold criteria

Venn diagrams were made according to the numerical size of interactors using the Venn diagram plotter (https://omics.pnl.gov/software/venn-diagram-plotter) developed by the Pacific Northwest National Laboratory (US Department of Energy). Two- and three-dimensional graphs were generated in Microsoft Excel and imported into Abobe Illustrator, where figures were assembled.

## Results and Discussion

### The interactome of SEPT9 isoforms i1, i4 and i5 reveal unique and overlapping binding partners

We sought to determine the interactome of SEPT9 isoforms 1 (SEPT9_i1), 4 (SEPT9_i4) and 5 (SEPT9_i5), which share the least overlap in their NTEs from all SEPT9 isoforms (Figure 1A) and therefore, may have differential interactions and functions. MCF-7 cell lines that stably express GFP-SEPT9_i1, GFP-SEPT9_i4 and GFP-SEPT9_i5 were previously established in order to investigate isoform-specific oncogenic properties (41). Owing to the low copy number of their endogenous SEPT9 gene (41), MCF-7 cells allow to recapitulate the upregulation of *SEPT9* isoforms that occurs in breast cancer cells by expressing exogenous GFP-tagged chimeras of *SEPT9* isoforms (Figure 2B). GFP-SEPT9 isoforms assembled into filaments (Figure 1B) and colocalized with actin filaments and/or microtubules (Supplementary Information, Figure S1), indicating that over-expression was not at levels that would disrupt assembly of SEPT9 isoforms into higher-order filaments and their interaction with the cytoskeleton.

To identify the interactome of each SEPT9 isoform, MCF-7 cells were lysed and immunoprecipitations were performed with an antibody against GFP coupled to an amine-reactive resin (Figure 1C). Lysates from untransfected MCF-7 cells were used as a negative control for identifying proteins that were precipitated non-specifically by the anti-GFP-coupled resin. Five independent immunoprecipitations were performed with lysates from each MCF-7 SEPT9 isoform-specific cell line. Protein-complexes were eluted from the antibody-bound resin and digested with trypsin prior to injecting into an ultra high-performance liquid chromatography (UHPLC) unit coupled to an electrospray ionization mass spectrometer (ESI-MS). Mass spectra were extracted and deconvoluted with the PEAKS Bioinformatics Studio software, and protein identities were derived by searches against the UniprotKB database for *Homo sapiens.*

For each isoform, protein hits were subjected to a four-tier stringency for the elimination of false positives. First, only protein hits with P-value (probability of false identification) score of −10lgP ≥ 15 in peptide-to-spectrum and peptide-to-protein matching were retained (55). Second, proteins that were identified in untransfected (control) cell lysates were removed from the proteome of each SEPT9 isoform. Third, protein hits that were not identified in at least two independent immunoprecipitations were excluded. Fourth, we removed protein hits that are most frequently identified as contaminants per the contaminant repository for affinity purification (CRAPome) database; protein hits that exceeded the score of 150 in the CRAPome database (Supplementary Information, Table S3).

By applying these stringency criteria, the interactome of SEPT9 isoforms 1, 4 and 5 consisted of 18, 20 and 24 proteins, respectively (Figure 2A-B). Approximately, a third of the binding partners of each isoform have been previously reported (Figure 2A), which validates our approach and indicates a high-confidence proteome. Of note, we detected a number of proteins that were previously reported to interact with SEPT9 (Table I, light shaded rows) or septins that form a complex with SEPT9, but did not meet stringency cutoffs; these included interactors that belonged to families of proteins of known association with septins (Table I, dark shaded rows). Hence, the interactomes of SEPT9 isoforms 1, 4 and 5 may exclude some *bona fide* interactions due to high-stringency criteria; a list of all protein hits from each independent experiment is provided in Supplementary Information Table S1.

The majority of the interactome (~75% of total proteins) of each SEPT9 isoform comprised non-septin proteins and nearly half (40-44%) was exclusively isoform-specific; i.e., binding partners were not shared with any other SEPT9 isoform. However, several proteins were common among all three isoforms or between two specific isoforms (Figure 2A). All three isoforms interacted with septins SEPT2, SEPT7 and SEPT8, which is consistent with previous findings of SEPT9 being part of hetero-octameric complex with subunits from the SEPT2, SEPT7 and SEPT6 groups; SEPT8 belongs to the SEPT6 group (14, 15). In addition to these septins, all three isoforms interacted with the adenomatous polyposis coli (APC) and the STE20-like serine/threonine protein kinase (SLK), which have been previously reported as septin interactors (54, 56), as well as titin (TTN) and keratin 23. SEPT9 isoforms 1 and 5 had two additional common binding partners (myomegalin and periplakin), while SEPT9 isoforms 4 and 5 shared SEPT11, the transmembrane protein 43 (TMEM43), the FER1-like family member 6 (FER1L6) and β-mannosidase as additional binding partners. Taken together, these data show that SEPT9 isoforms have common and unique interactors, which may arise respectively from shared protein domains (e.g., the GTP-binding domain) and differences in their NTEs.

### SEPT9 isoforms with shorter NTEs exhibit more promiscuous interactions with septins of the SEPT6 group

Septins were the most abundant group of proteins that co-immunoprecipitated with each SEPT9 isoform, which is consistent with their assembly into heteromeric complexes. SEPT9 itself was detected in all five independent immunoprecipitations against each isoform, indicating that SEPT9 was successfully immunoprecipitated in every run (Figure 3A). Collectively, the number of SEPT9 peptides were the most abundant - 56 for GFP-SEPT9_i1, 40 for GFP-SEPT9_i4 and 32 for GFP-SEPT9_i5 – and correlated with the increasingly shorter lengths of each SEPT9 isoform (Figure 3B).

**Figure 3.**
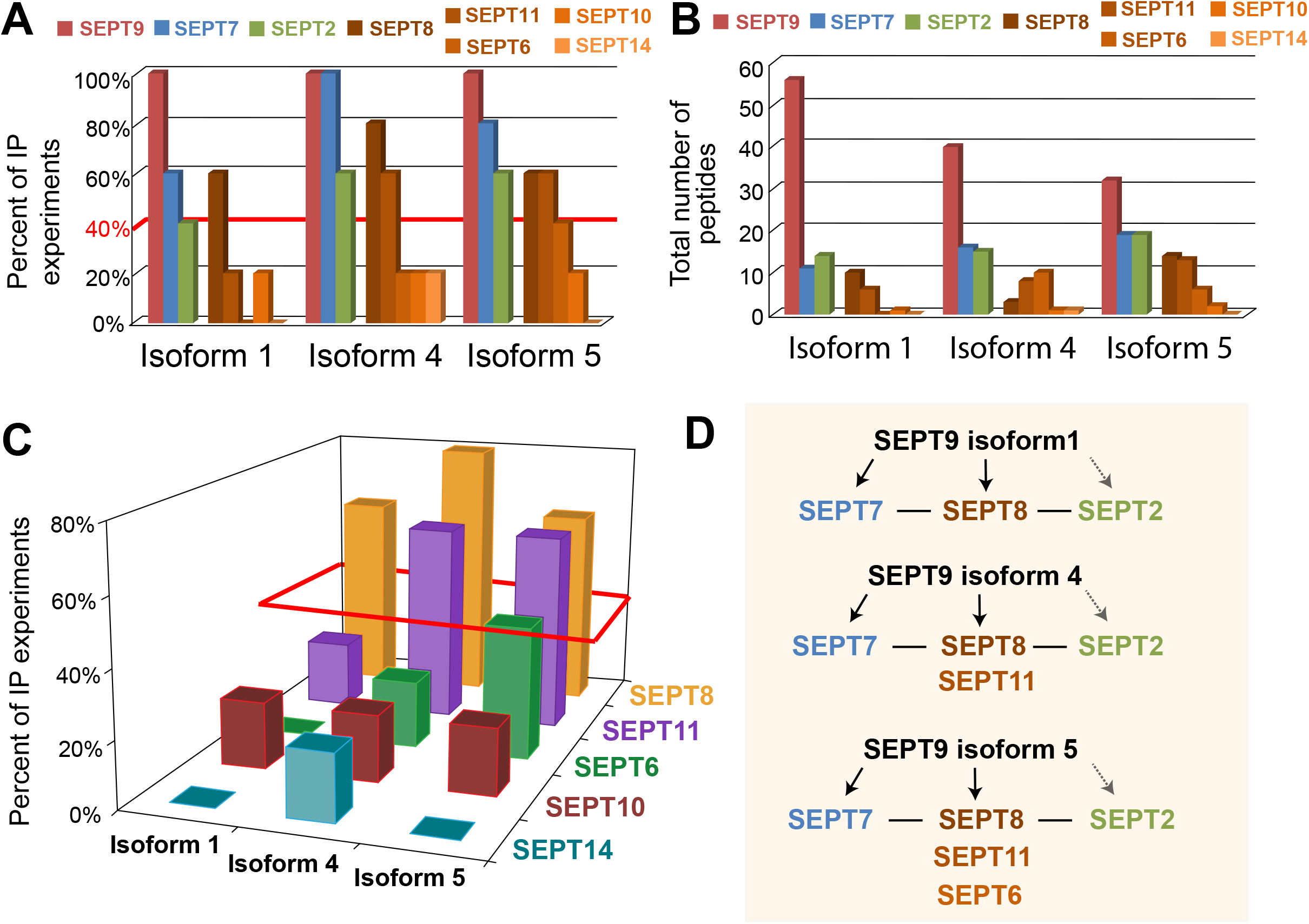
SEPT9 isoforms with shorter N-terminal extensions interact with multiple paralogs of the SEPT6 group. (A) Percent of IP experiments that resulted in the detection of septin paralogs (SEPT9, SEPT7, SEPT2, SEPT8, SEPT11, SEPT6, SEPT10, SEPT14) for each SEPT9 isoform. Red line indicates the cut-off (≥40% of experiments) for bona fide interactions. (B) Bar graph shows the cumulative number of septin peptides that were obtained from five independent immunoprecipitations of GFP-SEPT9 isoforms 1, 4 and 5. (C) Three-dimensional graph of the SEPT6 group paralogs (SEPT6, SEPT8, SEPT10, SEPT11, SEPT14) that co-IPed with SEPT9 isoforms 1, 4 and 5. The z axis indicates the percentage of experiments that resulted in detection of each SEPT6 group septin. The red parallelogram outlines the septin paralogs that were detected in ≥40% of the IP experiments. (D) Schematic summarizing the interactions of each SEPT9 isoform with septin paralogs. Based on previous evidence, solid arrows reflect direct interactions of SEPT9 with SEPT7 and potentially SEPT8, while the dashed opaque arrow indicates a potential indirect interaction with SEPT2 (via septins of the SEPT6 group). Solid lines denote direct interactions between SEPT7, SEPT6 group septins and SEPT2.

Among all septin paralogs that interacted with the SEPT9 isoforms, SEPT7 was most frequently pulled down (Figure 3A-B). Although less frequent than SEPT7, SEPT2 and SEPT8 were also present in the interactome of each SEPT9 isoform (Figure 3A-B). Strikingly, all other septin paralogs that co-immunoprecipitated with SEPT9 belonged to the SEPT6 group and their frequency of detection varied depending on SEPT9 isoform (Figure 3C). For SEPT9_i1, SEPT8 was the predominate paralog of the SEPT6 group; SEPT10 and SEPT11 were pulled down only once out of five immunoprecipitations. In contrast, SEPT9_i4 possessed both SEPT8 and SEPT11 as main binding partners of the SEPT6 group (Figure 3C). In immunoprecipitations of GFP-SEPT9_i5, SEPT6 emerged as a third major binding partner along with SEPT8 and SEPT11 (Figure 3A-C). These data reveal that SEPT9 isoforms with shorter NTEs have more promiscuous interactions with septins of the SEPT6 group.

Our results suggest that in MCF-7 breast cancer cells, SEPT9 isoforms form complexes with SEPT7, SEPT2 and septin paralogs of the SEPT6 group such as SEPT8, SEPT11 and SEPT6 (Figure 3D). This is consistent with previous studies showing that i) SEPT7 is a preferred and direct binding partner of SEPT9 (49, 50), and ii) members of the same septin group are exchangeable within the SEPT2/6/7/9 heteromer (13, 15). It is unclear, however, why there are no other members of SEPT2 group in complex with SEPT9 isoforms and why the truncated SEPT9 isoforms 4 and 5 allow for more flexibility in the exchange of SEPT6 group subunits than the full length SEPT9_i1. The former could be explained by low expression of other SEPT2 group septins in MCF-7 cells (57), while the latter suggests that SEPT9 isoforms possess or enable differential interactions with septin paralogs of the SEPT6 group. Interestingly, yeast two-hybrid screens have shown that septins of the SEPT6 group interact directly with SEPT9 (49). Moreover, yeast three-hybrid screens have demonstrated that SEPT9/6/7 complexes can be formed with SEPT9 taking the place of SEPT2 within the canonical SEPT2/6/7 complex (50). It is therefore plausible that SEPT9 isoforms hetero-dimerize with SEPT6 or form complexes with SEPT6/7 dimers. In this scenario, the full-length NTE of SEPT9_i1 could dictate preference for SEPT8, while the truncated NTEs of isoforms 4 and 5 allow for greater flexibility in interacting with SEPT 11 as well as SEPT6. In support of this possibility, the NTE of the yeast septin Cdc3 has been reported to influence binding to the septin paralogs Cdc10 versus Cdc12 through allosteric autoinhibitory interactions with the GTP-binding domain of Cdc3 (17). Hence, the lengths of the NTEs of SEPT9 isoforms could similarly impact binding to septins of the SEPT6 group, as N-terminal truncations could expose binding sites that allow for more promiscuous interactions; promiscuous septin-septin interactions have been shown to occur via the dimerization interface that involves the core GTP-binding domain (58).

### Spatial mapping of the SEPT9 interactome shows isoform-specific differences in binding partners of distinct subcellular localizations

To gain a better insight into the oncogenic pathways and properties of SEPT9 isoforms, we performed spatial and functional clustering of all non-septin binding partners. Based on the bioinformatics databases, we mapped the interactome of each SEPT9 isoform by assigning each protein to a subcellular organelle or structure; note that several proteins associate with more than a single organelle (Supplemental Information, Table S4). We used this categorization to quantify the relative distribution of the binding partners of SEPT9 isoforms in subcellular locales such as the cytoskeleton (actin filaments, microtubules, intermediate filaments), centrosomes, primary cilia, plasma membrane, nuclei, Golgi, endosomes/lysosomes, mitochondria and cell adhesions.

The spatial distributions of the interactomes of each SEPT9 isoform followed common patterns, but isoform specificities were also found in the organelle enrichment of binding partners (Figure 4A). Approximately half of the interactome of all three SEPT9 isoforms were proteins that localized to the cytoskeleton (20-30%) and the plasma membrane (15-20%). The majority of the cytoskeletal proteins (40-50%) associated with microtubules, while the remaining were linked to actin and intermediate filaments; only a single spectrin-associated protein (ankyrin 3) was found as an isoform 1-specific partner. The rest of the proteome exhibited a fairly equivalent distribution into the nucleus (5-15%), cell adhesions (~10%), the centrosome (5-10%), endolysosomes (3-8%), Golgi (~5%) and primary cilia (0-10%).

**Figure 4.**
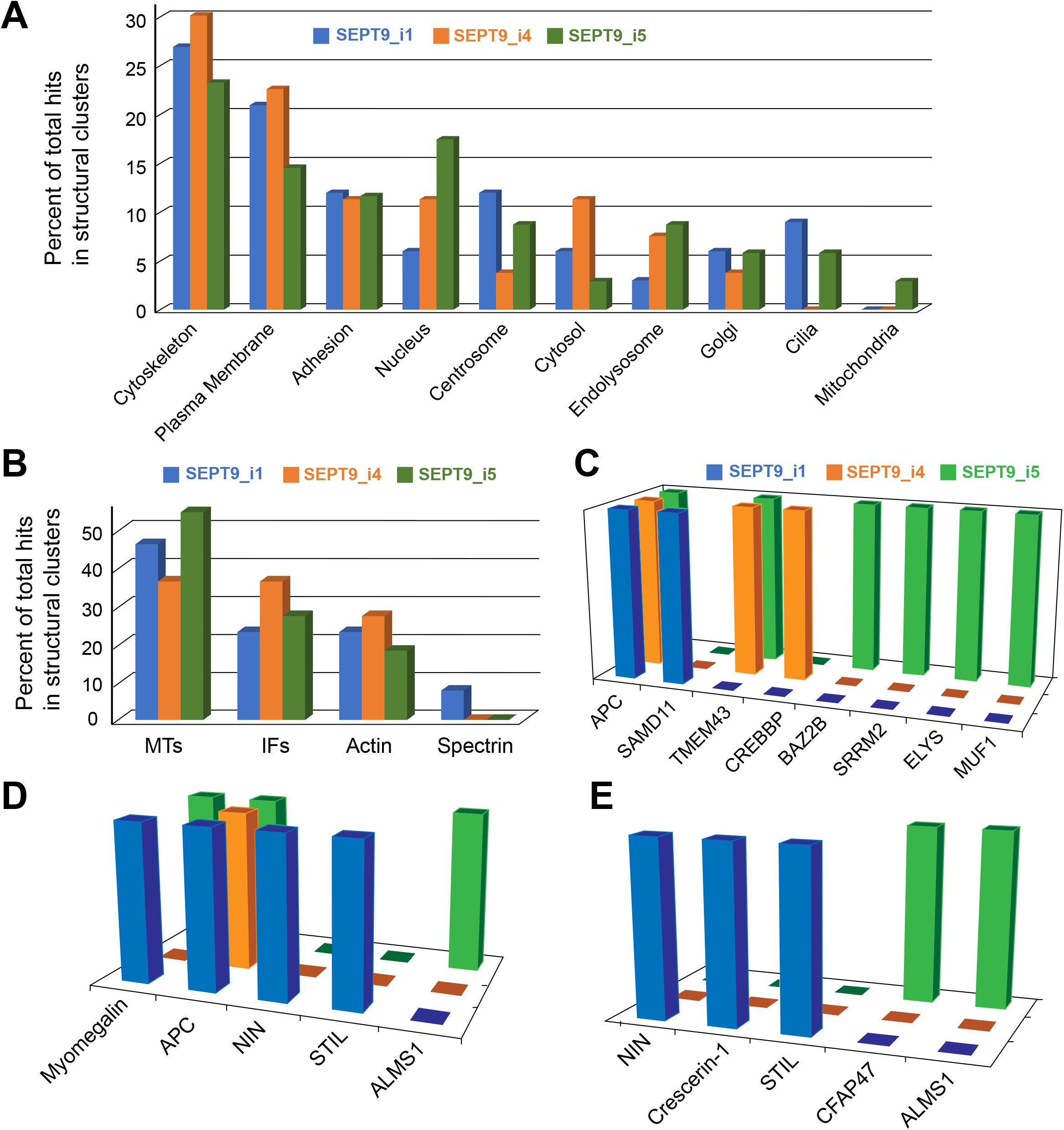
Spatial mapping of the SEPT9 interactome reveals isoform-specific interactions with proteins of distinct subcellular localizations. (A) The non-septin binding partners of SEPT9 isoforms 1, 4 and 5 were assigned to one or more subcellular locations based on published findings and bioinformatics databases. The protein number for each subcellular localization was divided by the total number of proteins in all subcellular clusters and plotted as percentage of total protein hits for each SEPT9 isoform. This provided an indication of the relative enrichment of the non-septin interactors of each SEPT9 isoform in distinct subcellular locales. (B) Protein interactors that were components of the MT, IF, actin and spectrin cytoskeletons or associated with these cytoskeletons were further subdivided according to their cytoskeletal identity and affiliation. Relative enrichment in each of the four cytoskeletal networks was quantified by calculating the percentage of total protein interactors assigned to each cytoskeleton. (C-E) Three-dimensional graphs show the identify of protein interactors with nuclear (C), centrosomal (D) and ciliary (E) localization for each SEPT9 isoform.

Relative enrichment of SEPT9 binding partners in the cytoskeleton, plasma membrane, Golgi or cell adhesions did not vary significantly across isoforms, but distinct differences were noted for proteins that localize to the nucleus, the centrosome and primary cilia. SEPT9 isoforms with truncated NTEs possessed a greater diversity of interactors with nuclear localization (Figure 4A and 4C), and several isoform-specific partners were revealed. More strikingly, there was a paucity of centrosomal and ciliary partners for SEPT9_i4, while isoforms 1 and 5 had unique and overlapping proteins that localized to the centrosome and primary cilia (Figure 4A and 4D-E).

### Differential interactions of SEPT9 isoforms with nuclear, centrosomal and ciliary proteins

Several oncogenes and tumor suppressors are nuclear proteins involved in gene transcription and DNA damage repair (59). Moreover, centrosomes and cilia are linked to cancer pathogenesis through their respective roles in genomic instability and signaling pathways that control cell growth and proliferation (e.g., hedgehog, notch and WNT) (60, 61). Therefore, SEPT9 isoform-specific interactions with nuclear, centrosomal and ciliary proteins could have implications for the mechanism of tumor progression, depending on the SEPT9 isoform whose expression is altered.

In the nuclear interactome of SEPT9, the tumor suppressor APC was a shared binding partner among all isoforms and TMEM43/LUMA was found as a common partner of isoforms 4 and 5; TMEM43/LUMA is an inner nuclear membrane protein that interfaces with epidermal growth factor receptor (EGFR) signaling pathways that control cell survival (62). The rest of the interactome was isoform specific (Figure 4C). Isoforms 1 and 4 interacted with SAMD11 (sterile alpha motif domain containing 11) and CREBBP (CREP-binding protein; CBP), respectively, while SEPT9_i4 pulled down BAZ2B (bromodomain adjacent to zinc finger domain 2B), SRRM2 (serine/arginine repetitive matrix protein 2), ELYS (embryonic large molecule derived from yolk sac; also known as AT-hook containing transcription factor 1) and MUF1 (also known as leucine-rich repeat containing protein 41, LRRC41).

Although little is known about SAMD11, its locus in hyper-methylated during breast cancer progression and is functionally implicated in transcriptional repression and cell proliferation (63). In contrast, CREBBP (CREB binding protein; CBP) is a well-studied transcriptional activator and histone acetyltransferase, which is posited to function as a scaffold or hub for various effectors of signaling pathways. Interestingly, BAZ2B is also a histone-binding protein with a bromodomain that recognizes acetylated lysines and might be involved in regulating the expression of noncoding RNAs (64). SRRM2 (SRM300) is involved in pre-mRNA splicing and verifiably, septins (SEPT9 included) were identified as interactors in a proteomic analysis of SRm160, which forms a complex with SRRM2 (65). ELYS/AHCTF1 is a component of the nuclear pore that interacts with chromatin (66). Recent studies show that ELYS is critical for nuclear pore assembly and coupling the inner nuclear membrane to heterochromatin through the lamin B receptor (67, 68). Together with BAZ2B and ELYS/AHCTF1, MUF1 is the third nuclear binding partner which was pulled down only with SEPT9_i5. MUF1 is not well characterized, but it is degraded in the cytoplasm by a RhoBTB-Cul3 ubiquitin ligase complex (69). Taken together, these interactions point to SEPT9 isoform-specific functions in epigenetic regulation of gene transcription, nucleo-cytoplasmic transport and nuclear envelope organization. Such roles are consistent with reports for SEPT9 localization to the nucleus (41) and data from transcriptomic profiling of MCF7 cells showing that SEPT9 isoforms have differential effects on gene expression (70), which are likely to arise due to differential interactions with transcription factors and histone modifiers. Additionally, the interactions of SEPT9_i5 with two nuclear membrane proteins (TMEM43, ELYS), which are involved in nuclear membrane interactions with nuclear lamins, suggest that SEPT9 isoforms may also impact gene expression through alteration of chromatin organization by nuclear membrane proteins and lamins (71, 72).

Similar to the nuclear interactome, several centrosomal and ciliary proteins were identified with specific SEPT9 isoforms (Figure 4D-E). A few of these proteins were common to centrosomes and cilia, which stem from centrioles and thus, have common protein components (73). Only two centrosomal proteins were shared across isoforms: APC, a multifunctional protein which also localizes to the centrosome (74), was identified as an interacting partner by all SEPT9 isoforms, and myomegalin was detected as a common binding partner of SEPT9_i1 and SEPT9_i5 (Figure 4D). Myomegalin synergizes with γ-tubulin and the end-binding protein 1 (EB1) for the nucleation of microtubules at the centrosome and Golgi membranes (75–78). Interestingly, recent studies indicate that SEPT1 is required for the nucleation of microtubules from Golgi membranes, suggesting that SEPT9 and myomegalin could be part of the underlying mechanism (79, 80). Strikingly, ninein and STIL (centriolar assembly protein) were identified as SEPT9_i1-specific partners, while ALMS1 was detected with only SEPT9_i5 (Figure 4D). Ninein is an essential component of the centrosome, which localizes to the subdistal appendages of centrioles and is required for microtubule nucleation and anchoring (81, 82). Conversely, STIL is essential for cell cycle-dependent biogenesis and duplication of centrioles, which is critical for the proper chromosome segregation during mitosis (83–86). Lastly, the SEPT9_i5 partner ALMS1 is required for centriole cohesion by regulating the localization of a protein (C-Nap1; centrosomal Nek2-associated protein 1), which functions in the organization of the interconnecting fibers that link the two centrioles together (87, 88). These data implicate SEPT9 isoforms 4 and 5 in distinct structural and functional properties of the centrosome.

Septins localize to the base and the axoneme of cilia, which originate from the conversion of a centriole into a basal body and its docking to the plasma membrane (73, 89, 90). Septins partake in the localization of ciliary proteins by supporting a diffusion barrier at the base of the cilia and regulate the length of the axoneme, but the underlying mechanisms are poorly understood (90, 91). Our data indicate that the SEPT9_i1 isoform associates with crescerin-1 and ninein, while the SEPT9_i4 isoform interacts with CFAP47 and ALMS1 (Figure 4E). The precise role of ninein in cilia is not understood, but it is required for cilia formation and thus, might be involved in the conversion of centrioles to basal bodies (73, 92). On the other hand, crescerin is involved in the regulation of ciliary length by functioning as a microtubule polymerase that adds tubulin subunits through its tubulin-binding TOG domains (93, 94). While the ciliary and flagella protein 47 (CFAP47) has not been studied, ALMS1 localizes to the basal bodies of primary cilia and is required for ciliary biogenesis, but its precise function is not understood (95, 96). These findings further highlight the known role of septins in ciliary biogenesis and length, and suggest that alterations in the expression of individual SEPT9 isoforms may impact ciliary structure and function in breast cancer cells.

### Functional clustering of the SEPT9 proteome points to isoform-specific roles in cell signaling and degradative pathways

Given the isoform-specific interactions with nuclear, centrosomal and ciliary proteins, we probed for isoform-specific functions by sorting the interactome of each SEPT9 isoform according to protein function. Based on published literature and bioinformatic databases, all non-septin binding partners were assigned to one or more functional clusters, which broadly represented the known functions of each protein interactor (Supplemental Information, Table S4). Cytoskeleton-related proteins were separated into components of cytoskeletal filaments, denoted as “cytoskeleton”, and regulators of the cytoskeletal organization, which were further grouped into microtubule-associated proteins (MAPs), actin-binding proteins (ABPs) and membrane-cytoskeleton adaptors. Functional clusters were also created based on protein involvement in cell signaling and adhesion, gene transcription/regulation, membrane fusion, post-translational modifications such as ubiquitination, phosphorylation (kinases) and N-glycan processing, regulation of small GTPases (GAPs/GEFs), mitophagy, apoptosis, mRNA splicing and nuclear envelope assembly. Each functional cluster contained one or more proteins, which were quantified as percentage of the total number of proteins in all clusters to indicate the relative enrichment of each cluster in the interactome of a SEPT9 isoform.

As predicted by the spatial organization of the interactome, cytoskeleton-related functions were the most enriched for all three isoforms (Figure 5A). The functional cluster of cytoskeletal regulation was highly enriched (~20-25% of the entire interactome), and the MAP, membrane-cytoskeleton adaptors and cell adhesion were also in the top five most-enriched clusters (Figure 5A). Notably, the interactome of the SEPT9 isoform 1 was notably more enriched in cytoskeletal regulators and MAPs compared to isoforms 4 and 5, which lack the N-terminal basic domain of SEPT9_i1 that interacts with microtubules and actin (25, 26).

**Figure 5.**
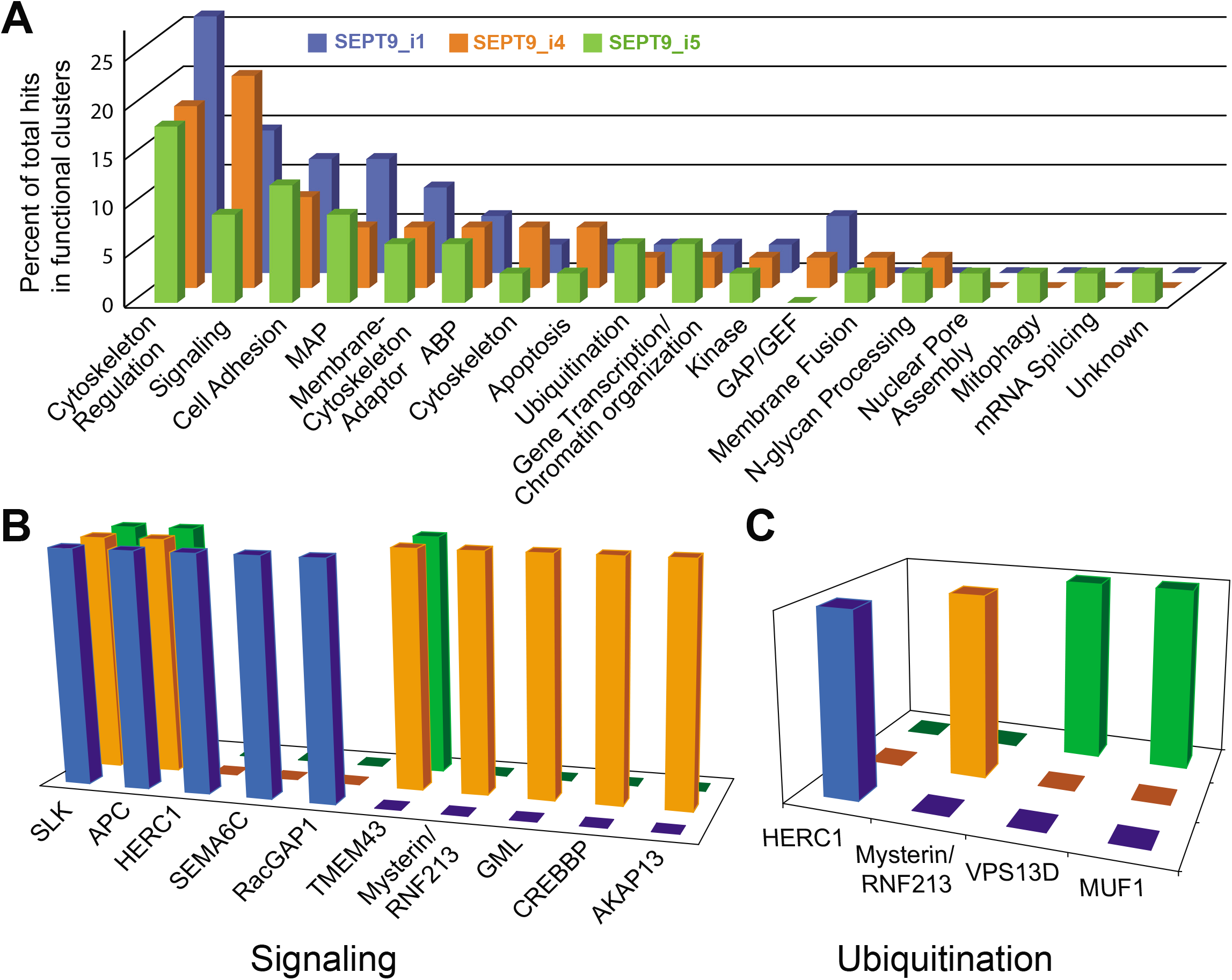
Functional clustering of the non-septin binding partners of SEPT9_i1, SEPT9_i4 and SEPT9_i5, and isoform-specific interactions with signaling and ubiquitinating factors. (A) Three-dimensional plot of the distribution of non-septin interactors of SEPT9_i1, SEPT9_i4 and SEPT9_i5 into functional clusters. Proteins of the interactome of each SEPT9 isoform were assigned into biological functions according to bioinformatic databases and published literature. Percent enrichment of each functional cluster was calculated by dividing the number of proteins in each cluster by the total number of proteins in all clusters, and multiplying by 100. (B-C) Three-dimensional graphs show the identify of protein interactors with signaling (B) and ubiquitinating (C) functions.

Strikingly, cell signaling was the most enriched cluster for SEPT9_i4 (Figure 5A). Compared to isoforms 1 and 5, the cell signaling cluster of SEPT9_i1 was 1.5- and 2.4-fold more enriched, respectively. The interactomes of isoforms 4 and 5 were more functionally diverse than isoform 1. In particular, isoform 5 exhibited the highest diversity, possessing a number of unique binding partners with functions in mRNA splicing, nuclear pore assembly and mitophagy. Overall, these isoform-specific differences correlate with protein length- and sequence-dependent differences such as: i) a cytoskeleton-binding domain in the N-terminus of SEPT9_i1, which appears to bias the interactome toward cytoskeleton-associated proteins; ii) a proline-rich domain in the N-terminal domain of SEPT9_i4, which in the absence of any upstream domains, could bias the interactome toward signaling proteins; proline-rich domains are common binding motifs for signaling molecules with SH3 domains; iii) lack of an NTE in the sequence of SEPT9_i5, which allows for more promiscuous interactions with septin paralogs of the SEPT6 group (Figure 3) and consequently, may result in more diverse interactions with non-septin proteins.

The elevated enrichment of the cell signaling cluster and the increased diversity of the SEPT9_i4 interactome bear significance in breast and ovarian cancers, in which SEPT9_i4 expression increases due to the expression of an mRNA transcript *(SEPT9_v4*)* with an alternative 5’-UTR sequence, which is translated more efficiently (97, 98). The cell signaling cluster of SEPT9_i4 contains four unique binding partners (Figure 5B): A-kinase anchoring protein 13 (AKAP13), the CREB-binding protein (CREBBP), GML (glycosylphospatidylinositol anchored molecule like) and mysterin/RNF213, all of which are linked to the pathology of breast cancer. AKAP13 is a RhoA GEF that activates Rho signaling and functions as a scaffold for the phosphorylation of the estrogen receptor alpha (ERα) by protein kinase A, which results in resistance to tamoxifen, an estrogen analog that targets ER positive breast tumors (99–101). Hence, SEPT9_i4 over-expression may impact both the pathogenesis and treatment of breast cancer through AKAP13. Conversely, CREBBP is a transcriptional activator that interacts with the BRCA1 tumor suppressor (102), which is the most frequently mutated gene in familial cases of breast cancer. GML is a target of the p53 tumor suppressor and it has been linked to apoptotic pathways that sensitize cancer cells to therapeutic treatments (103–105). Lastly, mysterin/RNF213 is a AAA ATPase with a ubiquitin ligase domain, which is implicated in signaling pathways involved in vascular development that supports cancer growth and metastasis (106, 107). Interestingly, mysterin is also involved in the stabilization of lipid droplets, which are regulated by SEPT9 in cells that are infected by the hepatitis C virus (108, 109).

In addition to the signaling cluster, ubiquitination also emerged as a distinct functional cluster with isoform-specific interactors (Figure 5C). SEPT9_i1 associated with HERC1, an E3 ubiqutin ligase that degrades the proto-oncogene serine/threonine-protein kinase c-RAF (110). The SEPT9_i4 interactome included mysterin/RNF213, which is an E3 ubiquitin ligase (107), and SEPT9_i5 pulled down VPS13D and MUF1. VPS13D has a ubiquitin-associated domain and is involved in mitochondrial clearance (mitophagy) (111), while MUF1 co-assembles with the E3 ubiquitin ligase Cullin/Rbx1 (112). The interaction with Vps13D is particularly interesting in the context of mitophagy as the mitochondrial SEPT5_i2 interacts with the E3 ubiquitin ligase parkin (113), and septins were recently shown to affect mitochondrial fission (114, 115). Collectively, these data implicate septins in ubiqutinase-based mechanisms of degradation and may inform future work on understanding how septins are degraded and/or regulate protein degradation. Of note, septins have been shown to inhibit the degradation of several proteins with roles in breast cancer including the EGF receptor, the Erb receptor tyrosine kinase 2 (ErbB2), the c-Jun-N-terminal kinase (JNK) and the hypoxia-inducible factor HIF1α, but the underlying mechanisms are not well understood (45, 116–118).

## Conclusion

The evolutionary expansion of the mammalian family of septins into a multitude of septin paralogs and isoforms suggests a functional specialization, which may arise from distinct interactions and protein-binding properties. However, this has yet to be explored as septin interactomes remain poorly characterized. Our results provide the first proteomic evidence of septin isoforms with differential interactions. Proteomic profiling of SEPT9 isoforms, which are over-expressed in breast cancer, revealed isoform-specific differences in interactions with septin paralogs and non-septin proteins of distinct subcellular localizations and functions. In agreement with recent findings in the heteromeric assembly of yeast septins (17), our data suggest that the NTEs of SEPT9 may allosterically determine the identity of their septin partners, which become more diverse with truncated N-terminal sequences. Similarly, we posit that the NTEs of SEPT9 isoforms impact interactions with non-septin proteins in a sequence- and length-dependent manner. Hence, the altered expression of specific SEPT9 isoforms in breast cancers could impact pathogenetic mechanisms and properties. Overall, this proteomic study implicates SEPT9 isoforms in hitherto unknown processes and mechanisms, which can benefit future studies of the oncogenic properties of SEPT9 and the design of therapies that target SEPT9 isoforms.

## Acknowledgments

We thank Drs. Matthew T. Balmer and Arun Arunachalam for their support and Sanofi Pasteur for use of the LCMS instrumentation and bioinformatics software. This work was supported with NIH/NIGMS grant RO1 GM097664 and a PA Department of Health CURE grant SAP 4100079710 to E.T.S. All microscopy took place in the Cell Imaging Center of Drexel University.

## Author Contributions

L.D. performed experiments, analyzed data, made figures and co-wrote manuscript with E.T.S.; G. P. contributed technically and intellectually to ESI/LC-MS/MS experimentation and software-based peptide identification; J.R.B. performed microscopy imaging; C.M. created MCF-7 cell lines and contributed to the manuscript; E.T.S. conceived and directed the project (experiments and analyses), drafted manuscript and edited figures.

## Supporting Information

Supporting Figures

Figure S1: Localization of GFP-SEPT9 isoforms 1, 4 and 5 with respect to the actin and microtubule cytoskeleton.

Supporting Tables

Table S1: List of proteins identified in all experimental runs

Table S2: Data for protein hits identified based on single peptides

Table S3: CRAPome frequencies of detection and spectral counts for all non-septin hits

Table S4: Spatial and functional categorization of all non-septin hits

**Figure S1:**
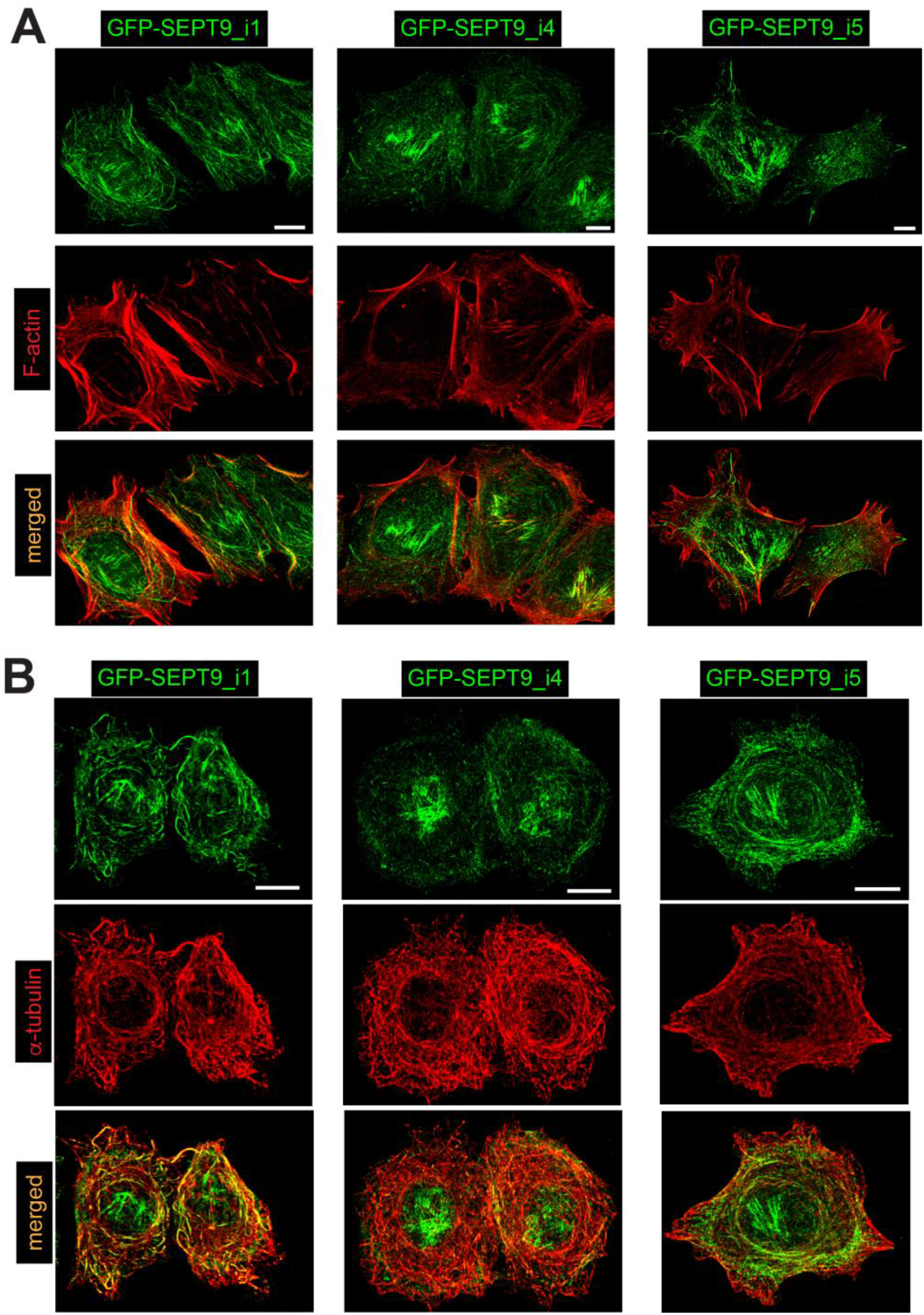
Localization of GFP-SEPT9 isoforms 1, 4 and 5 with respect to the actin and microtubule cytoskeleton. Images show maximal intensity projections of confocal microscopy sections taken from MCF-7 cells that express GFP-SEPT9_i1, GFP-SEPT9_i4 and GFP-SEPT9-i5 after staining for actin filaments with phalloidin (A) and microtubules with an antibody against α-tubulin (B). Scale bars, 10 μM.

**Supporting Tables**

**Table S1: List of proteins identified in all experimental runs.** A comprehensive list of all protein identities (protein ID, accession numbers, description) and their corresponding probabilities of false discovery (−10lgP score), percent coverage, number and uniqueness of peptides, modifications and average mass. Each sheet shows proteins identified in individual experimental runs (Runs 1 to 5) from five independent immunoprecipitations with lysates of MCF-7 cells expressing SEPT9_i1, SEPT9_i4 or SEPT9_i5, and untransfected control cells. A cut-off score of −10lgP ≥ 15 (false discovery probability in peptide-to-spectrum and peptide-to-protein matching) was applied.

**Table S2: Data for protein hits identified based on single peptides.** Sheet entitled “summary” shows all protein hits across five independent immunoprecipitations denoted as runs 1, 2, 3, 4 and 5. Numbers under each run correspond to the number of peptides detected and number of total peptides across all five runs is shown under column G labeled as “total”. Accession number (protein identity), amino acid sequence, spectral match number and −10lgP score, retention times and uniqueness (Y, yes; N, no) are shown for each single peptide. Sheets entitled Run 1, Run 2, Run 3, Run 4 and Run 5 include annotated MS/MS spectra with ion match table including b and y ions, and an error map (including error tolerance) for each identified peptide. The peptide spectrum match (PSM) view shows peptide sequence, precursor mass, mass to charge ratio (m/z), retention time, mass error (ppm) and −10lgP values.

**Table S3: CRAPome frequencies of detection and spectral counts for all non-septin hits.** According the CRAPome database, frequency of detection (number of experiments found out of total experiments), CRAPome score, percentage of times detected, average spectral counts (Ave SC) and maximal spectral counts (Max SC) are tabulated for each non-septin interactor of GFP-SEPT9_i1 (sheet 1), GFP-SEPT9_i4 (sheet 2) and GFP-SEPT9_i5 (sheet 3) that was not present in the negative control (untransfected MCF-7 cells) and met reproducibility, peptide-spectrum and peptide-protein threshold criteria. Shaded rows outline protein hits with CRAPome scores of >150, which were removed from the final list of protein hits.

**Table S4: Spatial and functional categorization of all non-septin hits.** Sheets 1 through 3 – All non-septin protein interactors of GFP-SEPT9_i1 (sheet 1), GFP-SEPT9_i4 (sheet 2) and GFP-SEPT9_i5 (sheet 3) are binned under subcellular organelles, structures and cytoskeletal systems; sheets 4 through 6 – all non-septin protein interactors of GFP-SEPT9_i1 (sheet 4), GFP-SEPT9_i4 (sheet 5) and GFP-SEPT9_i5 (sheet 6) are binned under subcellular organelles, structures and cytoskeletal systems.

## References

1. Mostowy, S.; Cossart, P., Septins: the fourth component of the cytoskeleton. Nat Rev Mol Cell Biol 2012, 13, (3), 183–94.

2. Kinoshita, M., Diversity of septin scaffolds. Curr Opin Cell Biol 2006, 18, (1), 54–60.

3. Gladfelter, A. S.; Pringle, J. R.; Lew, D. J., The septin cortex at the yeast mother-bud neck. Curr Opin Microbiol 2001, 4, (6), 681–9.

4. Caudron, F.; Barral, Y., Septins and the lateral compartmentalization of eukaryotic membranes. Dev Cell 2009, 16, (4), 493–506.

5. Spiliotis, E. T., Spatial effects - site-specific regulation of actin and microtubule organization by septin GTPases. J Cell Sci 2018, 131, (1).

6. Sirajuddin, M.; Farkasovsky, M.; Hauer, F.; Kuhlmann, D.; Macara, I. G.; Weyand, M.; Stark, H.; Wittinghofer, A., Structural insight into filament formation by mammalian septins. Nature 2007, 449, (7160), 311–5.

7. Valadares, N. F.; d’ Muniz Pereira, H.; Ulian Araujo, A. P.; Garratt, R. C., Septin structure and filament assembly. Biophys Rev 2017, 9, (5), 481–500.

8. Bridges, A. A.; Gladfelter, A. S., Septin Form and Function at the Cell Cortex. J Biol Chem 2015, 290, (28), 17173–80.

9. Dolat, L.; Hu, Q.; Spiliotis, E. T., Septin functions in organ system physiology and pathology. Biol Chem 2014, 395, (2), 123–41.

10. Kinoshita, M., The septins. Genome Biol 2003, 4, (11), 236.

11. Pan, F.; Malmberg, R. L.; Momany, M., Analysis of septins across kingdoms reveals orthology and new motifs. BMC Evol Biol 2007, 7, 103.

12. Russell, S. E.; Hall, P. A., Septin genomics: a road less travelled. Biol Chem 2011, 392, (8–9), 763–7.

13. Kinoshita, M., Assembly of mammalian septins. J Biochem 2003, 134, (4), 491–6.

14. Kim, M. S.; Froese, C. D.; Estey, M. P.; Trimble, W. S., SEPT9 occupies the terminal positions in septin octamers and mediates polymerization-dependent functions in abscission. J Cell Biol 2011, 195, (5), 815–26.

15. Sellin, M. E.; Sandblad, L.; Stenmark, S.; Gullberg, M., Deciphering the rules governing assembly order of mammalian septin complexes. Mol Biol Cell 2011, 22, (17), 3152–64.

16. Abbey, M.; Gaestel, M.; Menon, M. B., Septins: Active GTPases or just GTP-binding proteins? Cytoskeleton (Hoboken) 2018.

17. Weems, A.; McMurray, M., The step-wise pathway of septin hetero-octamer assembly in budding yeast. Elife 2017, 6.

18. Zent, E.; Wittinghofer, A., Human septin isoforms and the GDP-GTP cycle. Biol Chem 2014, 395, (2), 169–80.

19. Sellin, M. E.; Stenmark, S.; Gullberg, M., Cell type-specific expression of SEPT3-homology subgroup members controls the subunit number of heteromeric septin complexes. Mol Biol Cell 2014, 25, (10), 1594–607.

20. Sellin, M. E.; Stenmark, S.; Gullberg, M., Mammalian SEPT9 isoforms direct microtubule-dependent arrangements of septin core heteromers. Mol Biol Cell 2012, 23, (21), 4242–55.

21. Garcia, G., 3rd; Finnigan, G. C.; Heasley, L. R.; Sterling, S. M.; Aggarwal, A.; Pearson, C. G.; Nogales, E.; McMurray, M. A.; Thorner, J., Assembly, molecular organization, and membrane-binding properties of development-specific septins. J Cell Biol 2016, 212, (5), 515-29.

22. Neubauer, K.; Zieger, B., The Mammalian Septin Interactome. Front Cell Dev Biol 2017, 5, 3.

23. Fuchtbauer, A.; Lassen, L. B.; Jensen, A. B.; Howard, J.; Quiroga Ade, S.; Warming, S.; Sorensen, A. B.; Pedersen, F. S.; Fuchtbauer, E. M., Septin9 is involved in septin filament formation and cellular stability. Biol Chem 2011, 392, (8–9), 769–77.

24. McIlhatton, M. A.; Burrows, J. F.; Donaghy, P. G.; Chanduloy, S.; Johnston, P. G.; Russell, S. E., Genomic organization, complex splicing pattern and expression of a human septin gene on chromosome 17q25.3. Oncogene 2001, 20, (41), 5930–9.

25. Bai, X.; Bowen, J. R.; Knox, T. K.; Zhou, K.; Pendziwiat, M.; Kuhlenbaumer, G.; Sindelar, C. V.; Spiliotis, E. T., Novel septin 9 repeat motifs altered in neuralgic amyotrophy bind and bundle microtubules. J Cell Biol 2013, 203, (6), 895–905.

26. Smith, C.; Dolat, L.; Angelis, D.; Forgacs, E.; Spiliotis, E. T.; Galkin, V. E., Septin 9 Exhibits Polymorphic Binding to F-Actin and Inhibits Myosin and Cofilin Activity. J Mol Biol 2015, 427, (20), 3273–84.

27. Karasmanis, E. P.; Phan, C. T.; Angelis, D.; Kesisova, I. A.; Hoogenraad, C. C.; McKenney, R. J.; Spiliotis, E. T., Polarity of Neuronal Membrane Traffic Requires Sorting of Kinesin Motor Cargo during Entry into Dendrites by a Microtubule-Associated Septin. Dev Cell 2018, 46, (2), 204–218 e7.

28. Estey, M. P.; Di Ciano-Oliveira, C.; Froese, C. D.; Bejide, M. T.; Trimble, W. S., Distinct roles of septins in cytokinesis: SEPT9 mediates midbody abscission. J Cell Biol 2010, 191, (4), 741–9.

29. Russell, S. E.; McIlhatton, M. A.; Burrows, J. F.; Donaghy, P. G.; Chanduloy, S.; Petty, E. M.; Kalikin, L. M.; Church, S. W.; McIlroy, S.; Harkin, D. P.; Keilty, G. W.; Cranston, A. N.; Weissenbach, J.; Hickey, I.; Johnston, P. G., Isolation and mapping of a human septin gene to a region on chromosome 17q, commonly deleted in sporadic epithelial ovarian tumors. Cancer Res 2000, 60, (17), 4729–34.

30. Kalikin, L. M.; Sims, H. L.; Petty, E. M., Genomic and expression analyses of alternatively spliced transcripts of the MLL septin-like fusion gene (MSF) that map to a 17q25 region of loss in breast and ovarian tumors. Genomics 2000, 63, (2), 165–72.

31. Osaka, M.; Rowley, J. D.; Zeleznik-Le, N. J., MSF (MLL septin-like fusion), a fusion partner gene of MLL, in a therapy-related acute myeloid leukemia with a t(11;17)(q23;q25). Proc Natl Acad Sci U S A 1999, 96, (11), 6428–33.

32. Sorensen, A. B.; Lund, A. H.; Ethelberg, S.; Copeland, N. G.; Jenkins, N. A.; Pedersen, F.S., Sint1, a common integration site in SL3-3-induced T-cell lymphomas, harbors a putative proto-oncogene with homology to the septin gene family. J Virol 2000, 74, (5), 2161–8.

33. Montagna, C.; Lyu, M. S.; Hunter, K.; Lukes, L.; Lowther, W.; Reppert, T.; Hissong, B.; Weaver, Z.; Ried, T., The Septin 9 (MSF) gene is amplified and overexpressed in mouse mammary gland adenocarcinomas and human breast cancer cell lines. Cancer Res 2003, 63, (9), 2179–87.

34. Connolly, D.; Hoang, H. G.; Adler, E.; Tazearslan, C.; Simmons, N.; Bernard, V. V.; Castaldi, M.; Oktay, M. H.; Montagna, C., Septin 9 amplification and isoform-specific expression in peritumoral and tumor breast tissue. Biol Chem 2014, 395, (2), 157–67.

35. Chacko, A. D.; McDade, S. S.; Chanduloy, S.; Church, S. W.; Kennedy, R.; Price, J.; Hall, P. A.; Russell, S. E., Expression of the SEPT9_i4 isoform confers resistance to microtubule-interacting drugs. Cell Oncol (Dordr) 2012, 35, (2), 85–93.

36. Froidevaux-Klipfel, L.; Poirier, F.; Boursier, C.; Crepin, R.; Pous, C.; Baudin, B.; Baillet, A., Modulation of septin and molecular motor recruitment in the microtubule environment of the Taxol-resistant human breast cancer cell line MDA-MB-231. Proteomics 2011, 11, (19), 3877–86.

37. Froidevaux-Klipfel, L.; Targa, B.; Cantaloube, I.; Ahmed-Zaid, H.; Pous, C.; Baillet, A., Septin cooperation with tubulin polyglutamylation contributes to cancer cell adaptation to taxanes. Oncotarget 2015, 6, (34), 36063–80.

38. Amir, S.; Mabjeesh, N. J., SEPT9_V1 protein expression is associated with human cancer cell resistance to microtubule-disrupting agents. Cancer Biol Ther 2007, 6, (12), 1926–31.

39. Grutzmann, R.; Molnar, B.; Pilarsky, C.; Habermann, J. K.; Schlag, P. M.; Saeger, H. D.; Miehlke, S.; Stolz, T.; Model, F.; Roblick, U. J.; Bruch, H. P.; Koch, R.; Liebenberg, V.; Devos, T.; Song, X.; Day, R. H.; Sledziewski, A. Z.; Lofton-Day, C., Sensitive detection of colorectal cancer in peripheral blood by septin 9 DNA methylation assay. PLoS One 2008, 3, (11), e3759.

40. Warren, J. D.; Xiong, W.; Bunker, A. M.; Vaughn, C. P.; Furtado, L. V.; Roberts, W. L.; Fang, J. C.; Samowitz, W. S.; Heichman, K. A., Septin 9 methylated DNA is a sensitive and specific blood test for colorectal cancer. BMC Med 2011, 9, 133.

41. Connolly, D.; Yang, Z.; Castaldi, M.; Simmons, N.; Oktay, M. H.; Coniglio, S.; Fazzari, M. J.; Verdier-Pinard, P.; Montagna, C., Septin 9 isoform expression, localization and epigenetic changes during human and mouse breast cancer progression. Breast Cancer Res 2011, 13, (4), R76.

42. Chacko, A. D.; Hyland, P. L.; McDade, S. S.; Hamilton, P. W.; Russell, S. H.; Hall, P. A., SEPT9_v4 expression induces morphological change, increased motility and disturbed polarity. J Pathol 2005, 206, (4), 458–65.

43. Verdier-Pinard, P.; Salaun, D.; Bouguenina, H.; Shimada, S.; Pophillat, M.; Audebert, S.; Agavnian, E.; Coslet, S.; Charafe-Jauffret, E.; Tachibana, T.; Badache, A., Septin 9_i2 is downregulated in tumors, impairs cancer cell migration and alters subnuclear actin filaments. Sci Rep 2017, 7, 44976.

44. Gilad, R.; Meir, K.; Stein, I.; German, L.; Pikarsky, E.; Mabjeesh, N. J., High SEPT9_i1 protein expression is associated with high-grade prostate cancers. PLoS One 2015, 10, (4), e0124251.

45. Gonzalez, M. E.; Makarova, O.; Peterson, E. A.; Privette, L. M.; Petty, E. M., Up-regulation of SEPT9_v1 stabilizes c-Jun-N-terminal kinase and contributes to its pro-proliferative activity in mammary epithelial cells. Cell Signal 2009, 21, (4), 477–87.

46. Cravatt, B. F.; Simon, G. M.; Yates, J. R., 3rd, The biological impact of mass-spectrometry-based proteomics. Nature 2007, 450, (7172), 991–1000.

47. Walther, T. C.; Mann, M., Mass spectrometry-based proteomics in cell biology. J Cell Biol 2010, 190, (4), 491–500.

48. Larance, M.; Lamond, A. I., Multidimensional proteomics for cell biology. Nat Rev Mol Cell Biol 2015, 16, (5), 269–80.

49. Nakahira, M.; Macedo, J. N.; Seraphim, T. V.; Cavalcante, N.; Souza, T. A.; Damalio, J. C.; Reyes, L. F.; Assmann, E. M.; Alborghetti, M. R.; Garratt, R. C.; Araujo, A. P.; Zanchin, N. I.; Barbosa, J. A.; Kobarg, J., A draft of the human septin interactome. PLoS One 2010, 5, (11), e13799.

50. Sandrock, K.; Bartsch, I.; Blaser, S.; Busse, A.; Busse, E.; Zieger, B., Characterization of human septin interactions. Biol Chem 2011, 392, (8–9), 751–61.

51. Mellacheruvu, D.; Wright, Z.; Couzens, A. L.; Lambert, J. P.; St-Denis, N. A.; Li, T.; Miteva, Y. V.; Hauri, S.; Sardiu, M. E.; Low, T. Y.; Halim, V. A.; Bagshaw, R. D.; Hubner, N. C.; Al-Hakim, A.; Bouchard, A.; Faubert, D.; Fermin, D.; Dunham, W. H.; Goudreault, M.; Lin, Z. Y.; Badillo, B. G.; Pawson, T.; Durocher, D.; Coulombe, B.; Aebersold, R.; Superti-Furga, G.; Colinge, J.; Heck, A. J.; Choi, H.; Gstaiger, M.; Mohammed, S.; Cristea, I. M.; Bennett, K. L.; Washburn, M. P.; Raught, B.; Ewing, R. M.; Gingras, A. C.; Nesvizhskii, A. I., The CRAPome: a contaminant repository for affinity purification-mass spectrometry data. Nat Methods 2013, 10, (8), 730–6.

52. Gupta, N.; Pevzner, P. A., False discovery rates of protein identifications: a strike against the two-peptide rule. J Proteome Res 2009, 8, (9), 4173–81.

53. Perez-Riverol, Y.; Csordas, A.; Bai, J.; Bernal-Llinares, M.; Hewapathirana, S.; Kundu, D. J.; Inuganti, A.; Griss, J.; Mayer, G.; Eisenacher, M.; Perez, E.; Uszkoreit, J.; Pfeuffer, J.; Sachsenberg, T.; Yilmaz, S.; Tiwary, S.; Cox, J.; Audain, E.; Walzer, M.; Jarnuczak, A. F.; Ternent, T.; Brazma, A.; Vizcaino, J. A., The PRIDE database and related tools and resources in 2019: improving support for quantification data. Nucleic Acids Res 2019, 47, (D1), D442–D450.

54. Vargas-Muniz, J. M.; Renshaw, H.; Waitt, G.; Soderblom, E. J.; Moseley, M. A.; Palmer, J. M.; Juvvadi, P. R.; Keller, N. P.; Steinbach, W. J., Caspofungin exposure alters the core septin AspB interactome of Aspergillus fumigatus. Biochem Biophys Res Commun 2017, 485, (2), 221–226.

55. Ma, B.; Zhang, K.; Hendrie, C.; Liang, C.; Li, M.; Doherty-Kirby, A.; Lajoie, G., PEAKS: powerful software for peptide de novo sequencing by tandem mass spectrometry. Rapid Commun Mass Spectrom 2003, 17, (20), 2337–42.

56. Bejide, M. T. Characterization of a novel interaction between septins and the adenomatous polyposis coli tumor suppressor. University of Toronto, Toronto, Canada, 2010.

57. Orre, L. M.; Vesterlund, M.; Pan, Y.; Arslan, T.; Zhu, Y.; Fernandez Woodbridge, A.; Frings, O.; Fredlund, E.; Lehtio, J., SubCellBarCode: Proteome-wide Mapping of Protein Localization and Relocalization. Mol Cell 2019, 73, (1), 166–182 e7.

58. Kim, M. S.; Froese, C. D.; Xie, H.; Trimble, W. S., Uncovering principles that control septin-septin interactions. J Biol Chem 2012, 287, (36), 30406–13.

59. Lee, E. Y.; Muller, W. J., Oncogenes and tumor suppressor genes. Cold Spring Harb Perspect Biol 2010, 2, (10), a003236.

60. Godinho, S. A.; Pellman, D., Causes and consequences of centrosome abnormalities in cancer. Philos Trans R Soc Lond B Biol Sci 2014, 369, (1650).

61. Eguether, T.; Hahne, M., Mixed signals from the cell’s antennae: primary cilia in cancer. EMBO Rep 2018, 19, (11).

62. Jiang, H.; Hua, D.; Zhang, J.; Lan, Q.; Huang, Q.; Yoon, J. G.; Han, X.; Li, L.; Foltz, G.; Zheng, S.; Lin, B., MicroRNA-127-3p promotes glioblastoma cell migration and invasion by targeting the tumor-suppressor gene SEPT7. Oncol Rep 2014, 31, (5), 2261–9.

63. Titus, A. J.; Way, G. P.; Johnson, K. C.; Christensen, B. C., Deconvolution of DNA methylation identifies differentially methylated gene regions on 1p36 across breast cancer subtypes. Sci Rep 2017, 7, (1), 11594.

64. Kostrhon, S.; Kontaxis, G.; Kaufmann, T.; Schirghuber, E.; Kubicek, S.; Konrat, R.; Slade, D., A histone-mimicking interdomain linker in a multidomain protein modulates multivalent histone binding. J Biol Chem 2017, 292, (43), 17643–17657.

65. McCracken, S.; Longman, D.; Marcon, E.; Moens, P.; Downey, M.; Nickerson, J. A.; Jessberger, R.; Wilde, A.; Caceres, J. F.; Emili, A.; Blencowe, B. J., Proteomic analysis of SRm160-containing complexes reveals a conserved association with cohesin. J Biol Chem 2005, 280, (51), 42227–36.

66. Gillespie, P. J.; Khoudoli, G. A.; Stewart, G.; Swedlow, J. R.; Blow, J. J., ELYS/MEL-28 chromatin association coordinates nuclear pore complex assembly and replication licensing. Curr Biol 2007, 17, (19), 1657–62.

67. Mimura, Y.; Takagi, M.; Clever, M.; Imamoto, N., ELYS regulates the localization of LBR by modulating its phosphorylation state. J Cell Sci 2016, 129, (22), 4200–4212.

68. Clever, M.; Funakoshi, T.; Mimura, Y.; Takagi, M.; Imamoto, N., The nucleoporin ELYS/Mel28 regulates nuclear envelope subdomain formation in HeLa cells. Nucleus 2012, 3, (2), 187–99.

69. Schenkova, K.; Lutz, J.; Kopp, M.; Ramos, S.; Rivero, F., MUF1/leucine-rich repeat containing 41 (LRRC41), a substrate of RhoBTB-dependent cullin 3 ubiquitin ligase complexes, is a predominantly nuclear dimeric protein. J Mol Biol 2012, 422, (5), 659–673.

70. Zechmeister-Marcus, J.; Bejerano-Sagie, M.; Bagchi, S.; Verkhusha, V. V.; Connolly, D.; Goldberg, G. L.; Golden, A.; Sharma, V. P.; Condeelis, J.; Montagna, C., Septin 9 isoforms promote tumorigenesis in mammary epithelial cells by increasing migration and ECM degradation through metalloproteinase secretion at focal adhesions. (unpublished work, Albert Einstein College of Medicine) 2019.

71. Amendola, M.; van Steensel, B., Mechanisms and dynamics of nuclear lamina-genome interactions. Curr Opin Cell Biol 2014, 28, 61–8.

72. Raices, M.; D’Angelo, M. A., Nuclear pore complexes and regulation of gene expression. Curr Opin Cell Biol 2017, 46, 26–32.

73. Kobayashi, T.; Dynlacht, B. D., Regulating the transition from centriole to basal body. J Cell Biol 2011, 193, (3), 435–44.

74. Lui, C.; Mills, K.; Brocardo, M. G.; Sharma, M.; Henderson, B. R., APC as a mobile scaffold: regulation and function at the nucleus, centrosomes, and mitochondria. IUBMB Life 2012, 64, (3), 209–14.

75. Bouguenina, H.; Salaun, D.; Mangon, A.; Muller, L.; Baudelet, E.; Camoin, L.; Tachibana, T.; Cianferani, S.; Audebert, S.; Verdier-Pinard, P.; Badache, A., EB1-binding-myomegalin protein complex promotes centrosomal microtubules functions. Proc Natl Acad Sci US A 2017, 114, (50), E10687–E10696.

76. Roubin, R.; Acquaviva, C.; Chevrier, V.; Sedjai, F.; Zyss, D.; Birnbaum, D.; Rosnet, O., Myomegalin is necessary for the formation of centrosomal and Golgi-derived microtubules. Biol Open 2013, 2, (2), 238–50.

77. Verde, I.; Pahlke, G.; Salanova, M.; Zhang, G.; Wang, S.; Coletti, D.; Onuffer, J.; Jin, S. L.; Conti, M., Myomegalin is a novel protein of the golgi/centrosome that interacts with a cyclic nucleotide phosphodiesterase. J Biol Chem 2001, 276, (14), 11189–98.

78. Wang, Z.; Zhang, C.; Qi, R. Z., A newly identified myomegalin isoform functions in Golgi microtubule organization and ER-Golgi transport. J Cell Sci 2014, 127, (Pt 22), 4904–17.

79. Caudron, F.; Yadav, S., Meeting report - shining light on septins. J Cell Sci 2018, 131, (1).

80. Song, K.; Gras, C.; Capin, G.; Gimber, N.; Lehmann, M.; Mohd, S.; Puchkov, D.; Rodiger, M.; Wilhelmi, I.; Daumke, O.; Schmoranzer, J.; Schurmann, A.; Krauss, M., A SEPT1-based scaffold is required for Golgi integrity and function. J Cell Sci 2019, 132, (3).

81. Delgehyr, N.; Sillibourne, J.; Bornens, M., Microtubule nucleation and anchoring at the centrosome are independent processes linked by ninein function. J Cell Sci 2005, 118, (Pt 8), 1565–75.

82. Ou, Y. Y.; Mack, G. J.; Zhang, M.; Rattner, J. B., CEP110 and ninein are located in a specific domain of the centrosome associated with centrosome maturation. J Cell Sci 2002, 115, (Pt 9), 1825–35.

83. Arquint, C.; Sonnen, K. F.; Stierhof, Y. D.; Nigg, E. A., Cell-cycle-regulated expression of STIL controls centriole number in human cells. J Cell Sci 2012, 125, (Pt 5), 1342–52.

84. Moyer, T. C.; Clutario, K. M.; Lambrus, B. G.; Daggubati, V.; Holland, A. J., Binding of STIL to Plk4 activates kinase activity to promote centriole assembly. J Cell Biol 2015, 209, (6), 863–78.

85. Tang, C. J.; Lin, S. Y.; Hsu, W. B.; Lin, Y. N.; Wu, C. T.; Lin, Y. C.; Chang, C. W.; Wu, K. S.; Tang, T. K., The human microcephaly protein STIL interacts with CPAP and is required for procentriole formation. EMBO J 2011, 30, (23), 4790–804.

86. Vulprecht, J.; David, A.; Tibelius, A.; Castiel, A.; Konotop, G.; Liu, F.; Bestvater, F.; Raab, M. S.; Zentgraf, H.; Izraeli, S.; Kramer, A., STIL is required for centriole duplication in human cells. J Cell Sci 2012, 125, (Pt 5), 1353–62.

87. Knorz, V. J.; Spalluto, C.; Lessard, M.; Purvis, T. L.; Adigun, F. F.; Collin, G. B.; Hanley, N. A.; Wilson, D. I.; Hearn, T., Centriolar association of ALMS1 and likely centrosomal functions of the ALMS motif-containing proteins C10orf90 and KIAA1731. Mol Biol Cell 2010, 21, (21), 3617–29.

88. Yang, J.; Adamian, M.; Li, T., Rootletin interacts with C-Nap1 and may function as a physical linker between the pair of centrioles/basal bodies in cells. Mol Biol Cell 2006, 17, (2), 1033–40.

89. Hu, J.; Bai, X.; Bowen, J. R.; Dolat, L.; Korobova, F.; Yu, W.; Baas, P. W.; Svitkina, T.; Gallo, G.; Spiliotis, E. T., Septin-driven coordination of actin and microtubule remodeling regulates the collateral branching of axons. Curr Biol 2012, 22, (12), 1109–15.

90. Ghossoub, R.; Hu, Q.; Failler, M.; Rouyez, M. C.; Spitzbarth, B.; Mostowy, S.; Wolfrum, U.; Saunier, S.; Cossart, P.; Jamesnelson, W.; Benmerah, A., Septins 2, 7 and 9 and MAP4 colocalize along the axoneme in the primary cilium and control ciliary length. J Cell Sci 2013, 126, (Pt 12), 2583–94.

91. Hu, Q.; Milenkovic, L.; Jin, H.; Scott, M. P.; Nachury, M. V.; Spiliotis, E. T.; Nelson, W. J., A septin diffusion barrier at the base of the primary cilium maintains ciliary membrane protein distribution. Science 2010, 329, (5990), 436–9.

92. Graser, S.; Stierhof, Y. D.; Lavoie, S. B.; Gassner, O. S.; Lamla, S.; Le Clech, M.; Nigg, E. A., Cep164, a novel centriole appendage protein required for primary cilium formation. J Cell Biol 2007, 179, (2), 321–30.

93. Das, A.; Dickinson, D. J.; Wood, C. C.; Goldstein, B.; Slep, K. C., Crescerin uses a TOG domain array to regulate microtubules in the primary cilium. Mol Biol Cell 2015, 26, (23), 4248–64.

94. Louka, P.; Vasudevan, K. K.; Guha, M.; Joachimiak, E.; Wloga, D.; Tomasi, R. F.; Baroud, C. N.; Dupuis-Williams, P.; Galati, D. F.; Pearson, C. G.; Rice, L. M.; Moresco, J. J.; Yates, J. R., 3rd; Jiang, Y. Y.; Lechtreck, K.; Dentler, W.; Gaertig, J., Proteins that control the geometry of microtubules at the ends of cilia. J Cell Biol 2018, 217, (12), 4298–4313.

95. Hearn, T.; Spalluto, C.; Phillips, V. J.; Renforth, G. L.; Copin, N.; Hanley, N. A.; Wilson, D. I., Subcellular localization of ALMS1 supports involvement of centrosome and basal body dysfunction in the pathogenesis of obesity, insulin resistance, and type 2 diabetes. Diabetes 2005, 54, (5), 1581–7.

96. Li, G.; Vega, R.; Nelms, K.; Gekakis, N.; Goodnow, C.; McNamara, P.; Wu, H.; Hong, N. A.; Glynne, R., A role for Alstrom syndrome protein, alms1, in kidney ciliogenesis and cellular quiescence. PLoS Genet 2007, 3, (1), e8.

97. McDade, S. S.; Hall, P. A.; Russell, S. E., Translational control of SEPT9 isoforms is perturbed in disease. Hum Mol Genet 2007, 16, (7), 742–52.

98. Scott, M.; McCluggage, W. G.; Hillan, K. J.; Hall, P. A.; Russell, S. E., Altered patterns of transcription of the septin gene, SEPT9, in ovarian tumorigenesis. Int J Cancer 2006, 118, (5), 1325–9.

99. Bentin Toaldo, C.; Alexi, X.; Beelen, K.; Kok, M.; Hauptmann, M.; Jansen, M.; Berns, E.; Neefjes, J.; Linn, S.; Michalides, R.; Zwart, W., Protein Kinase A-induced tamoxifen resistance is mediated by anchoring protein AKAP13. BMC Cancer 2015, 15, 588.

100. Sterpetti, P.; Hack, A. A.; Bashar, M. P.; Park, B.; Cheng, S. D.; Knoll, J. H.; Urano, T.; Feig, L. A.; Toksoz, D., Activation of the Lbc Rho exchange factor proto-oncogene by truncation of an extended C terminus that regulates transformation and targeting. Mol Cell Biol 1999, 19, (2), 1334–45.

101. Diviani, D.; Soderling, J.; Scott, J. D., AKAP-Lbc anchors protein kinase A and nucleates Galpha 12-selective Rho-mediated stress fiber formation. J Biol Chem 2001, 276, (47), 44247–57.

102. Pao, G. M.; Janknecht, R.; Ruffner, H.; Hunter, T.; Verma, I. M., CBP/p300 interact with and function as transcriptional coactivators of BRCA1. Proc Natl Acad Sci U S A 2000, 97, (3), 1020–5.

103. Kimura, Y.; Furuhata, T.; Shiratsuchi, T.; Nishimori, H.; Hirata, K.; Nakamura, Y.; Tokino, T., GML sensitizes cancer cells to Taxol by induction of apoptosis. Oncogene 1997, 15, (11), 1369–74.

104. Kagawa, K.; Inoue, T.; Tokino, T.; Nakamura, Y.; Akiyama, T., Overexpression of GML promotes radiation-induced cell cycle arrest and apoptosis. Biochem Biophys Res Commun 1997, 241, (2), 481–5.

105. Higashiyama, M.; Miyoshi, Y.; Kodama, K.; Yokouchi, H.; Takami, K.; Nishijima, M.; Nakayama, T.; Kobayashi, H.; Minamigawa, K.; Nakamura, Y., p53-regulated GML gene expression in non-small cell lung cancer. a promising relationship to cisplatin chemosensitivity. Eur J Cancer 2000, 36, (4), 489–95.

106. Kobayashi, H.; Matsuda, Y.; Hitomi, T.; Okuda, H.; Shioi, H.; Matsuda, T.; Imai, H.; Sone, M.; Taura, D.; Harada, K. H.; Habu, T.; Takagi, Y.; Miyamoto, S.; Koizumi, A., Biochemical and Functional Characterization of RNF213 (Mysterin) R4810K, a Susceptibility Mutation of Moyamoya Disease, in Angiogenesis In Vitro and In Vivo. J Am Heart Assoc 2015, 4, (7).

107. Liu, W.; Morito, D.; Takashima, S.; Mineharu, Y.; Kobayashi, H.; Hitomi, T.; Hashikata, H.; Matsuura, N.; Yamazaki, S.; Toyoda, A.; Kikuta, K.; Takagi, Y.; Harada, K. H.; Fujiyama, A.; Herzig, R.; Krischek, B.; Zou, L.; Kim, J. E.; Kitakaze, M.; Miyamoto, S.; Nagata, K.; Hashimoto, N.; Koizumi, A., Identification of RNF213 as a susceptibility gene for moyamoya disease and its possible role in vascular development. PLoS One 2011, 6, (7), e22542.

108. Akil, A.; Peng, J.; Omrane, M.; Gondeau, C.; Desterke, C.; Marin, M.; Tronchere, H.; Taveneau, C.; Sar, S.; Briolotti, P.; Benjelloun, S.; Benjouad, A.; Maurel, P.; Thiers, V.; Bressanelli, S.; Samuel, D.; Brechot, C.; Gassama-Diagne, A., Septin 9 induces lipid droplets growth by a phosphatidylinositol-5-phosphate and microtubule-dependent mechanism hijacked by HCV. Nat Commun 2016, 7, 12203.

109. Sugihara, M.; Morito, D.; Ainuki, S.; Hirano, Y.; Ogino, K.; Kitamura, A.; Hirata, H.; Nagata, K., The AAA+ ATPase/ubiquitin ligase mysterin stabilizes cytoplasmic lipid droplets. J Cell Biol 2019.

110. Schneider, T.; Martinez-Martinez, A.; Cubillos-Rojas, M.; Bartrons, R.; Ventura, F.; Rosa, J. L., The E3 ubiquitin ligase HERC1 controls the ERK signaling pathway targeting C-RAF for degradation. Oncotarget 2018, 9, (59), 31531–31548.

111. Anding, A. L.; Wang, C.; Chang, T. K.; Sliter, D. A.; Powers, C. M.; Hofmann, K.; Youle, R. J.; Baehrecke, E. H., Vps13D Encodes a Ubiquitin-Binding Protein that Is Required for the Regulation of Mitochondrial Size and Clearance. Curr Biol 2018, 28, (2), 287–295 e6.

112. Kamura, T.; Burian, D.; Yan, Q.; Schmidt, S. L.; Lane, W. S.; Querido, E.; Branton, P. E.; Shilatifard, A.; Conaway, R. C.; Conaway, J. W., Muf1, a novel Elongin BC-interacting leucine-rich repeat protein that can assemble with Cul5 and Rbx1 to reconstitute a ubiquitin ligase. J Biol Chem 2001, 276, (32), 29748–53.

113. Choi, P.; Snyder, H.; Petrucelli, L.; Theisler, C.; Chong, M.; Zhang, Y.; Lim, K.; Chung, K. K.; Kehoe, K.; D’Adamio, L.; Lee, J. M.; Cochran, E.; Bowser, R.; Dawson, T. M.; Wolozin, B., SEPT5_v2 is a parkin-binding protein. Brain Res Mol Brain Res 2003, 117, (2), 179–89.

114. Pagliuso, A.; Tham, T. N.; Stevens, J. K.; Lagache, T.; Persson, R.; Salles, A.; Olivo-Marin, J. C.; Oddos, S.; Spang, A.; Cossart, P.; Stavru, F., A role for septin 2 in Drp1-mediated mitochondrial fission. EMBO Rep 2016, 17, (6), 858–73.

115. Sirianni, A.; Krokowski, S.; Lobato-Marquez, D.; Buranyi, S.; Pfanzelter, J.; Galea, D.; Willis, A.; Culley, S.; Henriques, R.; Larrouy-Maumus, G.; Hollinshead, M.; Sancho-Shimizu, V.; Way, M.; Mostowy, S., Mitochondria mediate septin cage assembly to promote autophagy of Shigella. EMBO Rep 2016, 17, (7), 1029–43.

116. Amir, S.; Wang, R.; Simons, J. W.; Mabjeesh, N. J., SEPT9_v1 up-regulates hypoxia-inducible factor 1 by preventing its RACK1-mediated degradation. J Biol Chem 2009, 284, (17), 11142–51.

117. Marcus, E. A.; Tokhtaeva, E.; Turdikulova, S.; Capri, J.; Whitelegge, J. P.; Scott, D. R.; Sachs, G.; Berditchevski, F.; Vagin, O., Septin oligomerization regulates persistent expression of ErbB2/HER2 in gastric cancer cells. Biochem J 2016, 473, (12), 1703–18.

118. Diesenberg, K.; Beerbaum, M.; Fink, U.; Schmieder, P.; Krauss, M., SEPT9 negatively regulates ubiquitin-dependent downregulation of EGFR. J Cell Sci 2015, 128, (2), 397–407.

